# Human striatal progenitor cells that contain inducible safeguards and overexpress BDNF rescue Huntington’s disease phenotypes in R6/2 mice

**DOI:** 10.1101/2024.05.01.592095

**Authors:** Danielle A. Simmons, Sridhar Selvaraj, Tingshuo Chen, Gloria Cao, Talita Sauto Camelo, Tyne L.M. McHugh, Selena Gonzalez, Renata Martin, Juste Simanauskaite, Nobuko Uchida, Matthew Porteus, Frank M. Longo

## Abstract

Huntington’s disease (HD) is an autosomal-dominant neurodegenerative disorder characterized by striatal atrophy. Reduced trophic support due to decreased striatal levels of neurotrophins (NTs), mainly brain-derived neurotrophic factor (BDNF), contributes importantly to HD pathogenesis; restoring NTs has significant therapeutic potential. Human pluripotent stem cells (hPSC) offer a scalable platform for NT delivery but has potential safety risks including teratoma formation. We engineered hPSCs to constitutively produce BDNF and contain inducible safeguards to eliminate these cells if safety concerns arise. This study examined the efficacy of intrastriatally transplanted striatal progenitor cells (STRpcs) derived from these hPSCs against HD phenotypes in R6/2 mice. Engrafted STRpcs overexpressing BDNF alleviated motor and cognitive deficits and reduced mutant huntingtin aggregates. Activating the inducible safety switch with rapamycin safely eliminated the engrafted cells. These results demonstrate that BDNF delivery via a novel hPSC-based platform incorporating safety switches could be a safe and effective HD therapeutic.

## Introduction

Huntington’s Disease (HD) is an autosomal dominant neurodegenerative disorder caused by an expanded CAG repeat in the first exon of the *huntingtin* gene (*HTT*) encoding a mutant huntingtin (mHtt) protein with an extended stretch of polyglutamines^1^. mHtt aggregates in the cytoplasm and nucleus of cells and causes the preferential degeneration of medium spiny neurons (MSNs) in the striatum, eventually leading to atrophy of other brain regions, including the cortex^2^. This neurodegeneration underlies the progressive motor and cognitive deficits and psychiatric disturbances characteristic of HD. To date, no therapy exists that can delay the onset or slow the progression of HD and there is a critical need for disease-modifying therapies.

Many neurodegenerative mechanisms contribute to HD pathogenesis however an essential contributor is diminished neurotrophic support primarily due to the severe loss of brain-derived neurotrophic factor (BDNF) and signaling through its TrkB receptor^3,4^. BDNF is an essential trophic factor for striatal MSNs and cortical projection neurons, the earliest affected and most vulnerable cell types in HD brains^4,5^. Neurotrophin-3 (NT3) is another growth factor that has been shown to promote striatal synaptic plasticity and provide trophic support to motor neurons^6-8^ and cortical NT3 mRNA levels are greatly reduced in HD patients^9^. Infusing BDNF or NT3 into the striatum of rodent models of HD is neuroprotective^10,11^. Trophic factor deficits play a key role in HD-related neuropathology and many preclinical studies have shown that restoring BDNF or NT3 and their signaling may be an extremely effective therapeutic strategy against HD neurodegeneration^3,4,12^.

Cellular methods of NT delivery, particularly stem cell-based therapies, have produced promising therapeutic effects for many brain disorders and neurodegenerative diseases, including Parkinson’s disease and HD^13-15^. Stem cell-based therapies can potentially target multiple disease mechanisms by replacing degenerated cells and/or secreting neuroprotective factors, such as NTs, a large number of HD preclinical studies have used this approach^13,15-21^. Most of these studies utilized murine stem cells, typically mesenchymal stem cells, intracranially transplanted in HD rodent models and attributed the observed beneficial effects to trophic factors, primarily BDNF. Few studies utilized pluripotent stem cells (PSCs), including embryonic stem cells (ESCs) and induced pluripotent stem cells (iPSCs). PSCs offer features conducive to successful cell-based therapies such as unlimited expansion potential and the possibility of differentiating into all somatic cell types ^22^. However, PSC-based therapies are associated with possible safety risks including teratomas, which are a type of tumor that can form from residual undifferentiated PSCs in the therapeutic cell product ^23^. Another safety risk is *in vivo* engraftment of undesired cell types which could occur if the therapeutic cell product contains contaminating cells of a wrong lineage^24^. Thus, stem-cell based therapeutic strategies for HD and other neurodegenerative diseases would benefit greatly from using human PSCs (hPSCs) and incorporating safeguards to minimize or eliminate these potential hazards.

To address these safety risks, we previously reported engineering PSCs to express orthogonal dual safety switches using the highly efficient Cas9/guide RNA ribonucleoprotein (RNP) and AAV6-based gene editing platform. The safety switches were validated *in vivo* and shown to be efficacious in preventing teratomas and eliminating PSC-derived tumors when activated with orthogonal small molecules ^25^. Here, we used hPSCs expressing the safety switches to test the feasibility of developing an hPSC-based therapeutic platform to deliver NTs to the brains of an HD mouse model. Specifically, this proof-of-concept study was designed to determine whether intrastriatal transplantation of human neural progenitor cells (hNPCs) or striatal progenitor cells (hSTRpcs) derived from hPSCs containing orthogonal safeguards and engineered to overexpress either BDNF or NT3 can provide neuroprotection and prevent functional deficits in the R6/2 mouse model of HD. Moreover, we tested efficacy of the safety switches to remove the engrafted cells from the brain. This study provides *in vivo* evidence of the effectiveness of adding an engineered safeguard to an hPSC-based neuroregenerative therapeutic strategy for HD and other neurodegenerative diseases.

## RESULTS

### Generation of hPSC cell lines overexpressing BDNF and NT3 from hESCs with dual safety switches

hPSCs overexpressing BDNF or NT3 were derived from previously generated hESCs (line H9) expressing orthogonal, small molecule, inducible, dual safety switches^25^. The first safety switch is expressed under the control of endogenous NANOG promoter and is specifically expressed in PSCs. This safety switch transgene consists of Caspase9 (CASP9) fused to a mutant FKBP domain (F36V) and can be activated using a small molecule ligand, AP20187, to induce dimerization of the mutant FKBP domain (F36V) (**Supp. Fig. 1A**). The second safety switch is expressed under the control of the endogenous ACTB promoter and thus it is ubiquitously expressed in all cell types. This safety switch transgene consists of CASP9 fused to the mutant FRB (mFRB) and FKBP domains; these domains heterodimerize when activated by AP21967 or rapamycin (**Supp. Fig. 1B**) ^25^. Activating either switch triggers the formation of CASP9 dimers which induces apoptosis and cell death.

The current study used these dual safety switch expressing H9 hESCs for further engineering to overexpress either BDNF or NT3. For the overexpression of NTs, we performed gene targeting at the *HBB* locus using Cas9 RNP and AAV6-based gene editing. *HBB* is a safe harbor for neural cell types since it has no functional role in these cells and the locus is not adjacent to any known oncogenes or other genes that might affect neural cell behavior. We utilized an sgRNA targeting exon 1 of *HBB* gene that was used in a gene editing clinical trial for sickle cell disease ^26^. An AAV6 HDR donor vector was generated with homology arms flanking the sgRNA genomic target site and insert sequence consisting of BDNF or NT3 cDNA under the constitutive ubiquitin C promoter and bGH polyA sequence (**Fig. 1A**). We assessed the frequency of gene targeting at two different AAV6 multiplicities of infection (MOIs), 5K and 10K, using ddPCR analysis. We observed gene targeting frequencies of 63% and 67% at 5K and 10K MOI in bulk edited cells for NT3 expression (**Fig. 1B**).

**Figure 1.**
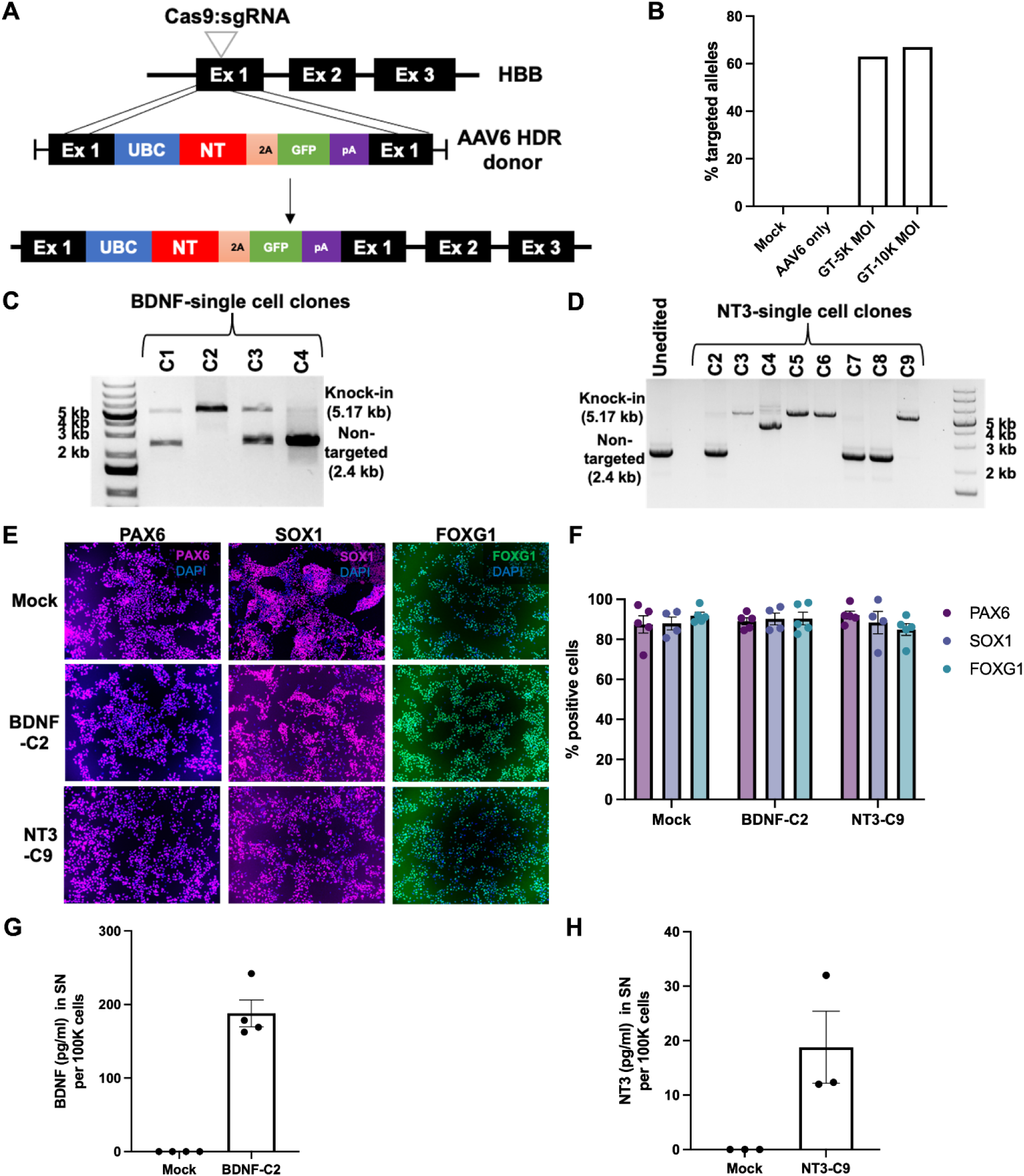
Generation of BDNF and NT3 overexpressing hPSCs and differentiation into hNPCs. (**A**) Gene targeting strategy at the *HBB* locus for generating hPSCs overexpressing BDNF or NT3 using CASP9 RNP and AAV6 gene editing. (**B**) Frequency of gene targeting at *HBB* locus for knock-in of NT3 construct post gene editing with AAV6 multiplicities of infection (MOIs) of 5K and 10K in bulk edited hESCs. GT denotes gene targeting with RNP and AAV6. (**C,D**) PCR for amplifying the region spanning knock-in of BDNF (**C**) and NT3 (**D**) overexpression constructs at the *HBB* locus in gene targeted single cell ESC clones. (**E**) Representative immunostaining images for neural progenitor cell (NPC) markers PAX6, SOX1, and the forebrain marker FOXG1 in hNPCs derived from mock, BDNF-C2, and NT3-C9 hPSCs. (**F**) Frequency of PAX6, SOX1, and FOXG1 positive NPCs derived from mock, BDNF-C2 and NT3-C9 hPSCs. Immunostaining images were quantified as area of corresponding marker staining relative to that of DAPI and represented as percentage of positive cells. (**G, H**) Bar graphs showing the quantification of ELISAs for BDNF (**G**) and NT3 (**H**) in the supernatants (SN) from hNPCs derived from BDNF-C2 and NT3-C9 hPSCs, respectively. Mock NPCs were used as a negative control. Data is shown as the concentration of BDNF or NT3 in the SN per 100K cells.

The gene-edited hPSCs expressing BDNF and NT3 were then subjected to single cell cloning to derive bi-allelic clonal cell lines. The isolated single cell clones were genotyped through PCR amplification of the region spanning knock-in. Genotyping of BDNF-expressing hPSC clones showed 1 bi-allelic knock-in (C2), 2 mono-allelic knock-in (C1 and C3), and 1 non-gene targeted (C4) clone (**Fig. 1C**). From NT3 clones, we derived 4 bi-allelic knock-in (C3, C5, C6 and C9) and 3 non-gene targeted (C2, C7, C8) clones (**Fig. 1D**). We chose one bi-allelic knock-in clone for BDNF (C2) and one for NT3 (C9) for further studies.

To confirm the expression and secretion of the NT factors, we performed ELISAs for BDNF and NT3 in the supernatants from the gene targeted BDNF and NT3 hPSC clones, respectively. Mean concentrations of 730 pg/ml BDNF and 153.5 pg/ml NT3 were determined per 100K cells for the BDNF-C2 and NT3-C9 clones, respectively (**Suppl. Fig. 2A, B**). We then tested whether overexpressing BDNF or NT3 affects pluripotency by assessing immunostaining for the pluripotency factors OCT3/4, SOX2 and NANOG. Both the BDNF and NT3 overexpressing hPSC lines showed robust expression of all three pluripotency markers and quantification confirmed that more than 95% of cells stained positive for the markers (**Supp. Fig. 2C, D**). These analyses showed that we generated hPSC lines that overexpress BDNF and NT3 and that these cell lines maintained their pluripotency marker expression.

### Derivation of neural progenitors from BDNF and NT3 overexpressing hPSCs

The BDNF- and NT3-overexpressing hPSCs were differentiated into hNPCs for *in vivo* transplantation studies. We used a previously described protocol involving small molecule-based dual SMAD inhibition (BMP and TGFβ inhibition) to induce differentiation into neuroectoderm. For differentiation into anterior neuroectoderm towards a forebrain fate, we also used a small molecule to inhibit WNT signaling ^27^. After 12 days of differentiation, we derived hNPCs from mock, BDNF-C2, and NT3-C9 hPSC lines. hNPCs robustly expressed the neural progenitor markers, PAX6 and SOX1, and the forebrain marker, FOXG1, based on immunostaining (**Fig. 1E**). Quantification of the immunostaining indicated that more than 80% of hNPCs from all three hPSC lines stained positive for PAX6, SOX1, and FOXG1 (**Fig. 1F**). To verify secretion of BDNF and NT3, we performed ELISA on the supernatants from the hNPCs. We detected the mean BDNF and NT3 concentrations to be approximately 188 pg/ml and 19 pg/ml per 100K cells, respectively, in the supernatants from the corresponding hNPCs (**Fig. 1G, H**). In supernatants of mock hNPCs, BDNF or NT3 were not detectable. BDNF and NT3 levels were maintained in the supernatants for hNPCs albeit at a lower level than was found in the supernatants of the undifferentiated hPSCs. Since, we detected NT secretion only in the hNPCs derived from gene targeted hPSCs, we proceeded with *in vivo* studies.

### Intrastriatally transplanted hNPCs overexpressing NT3 or BDNF prevent behavior deficits in R6/2 mice

The initial part of these experiments was designed to determine if intrastriatal transplantation of hNPCs engineered to overexpress BDNF or NT3 can prevent functional deficits in R6/2 mice compared to hNPCs without NT overexpression. After identifying the cell type that produced the most robust effects, our goal was to further mature these cells to striatal neuron progenitors (hSTRpcs) and assess them in a follow-up, statistically-powered preclinical efficacy study. This study used R6/2 mice which are transgenic for the 5′ end of the human HD gene carrying 100–190 glutamine (CAG) repeats and are a good model of juvenile HD or the anomalous splicing of mHtt that occurs in HD ^28^. They have rapidly developed, well-characterized pathological features including intranuclear mHtt aggregates developing at 4 weeks of age, progressive cognitive and motor deficits at 5–7 weeks, and a lifespan of ∼12 weeks ^28,29^.

The experimental timeline is illustrated in **Figure 2A**. Intrastriatal transplantation surgeries were performed on WT and R6/2 mice aged 4.5 to 5 weeks after the mice were administered an antibody-based immunosuppression protocol for 2 days (**Fig. 2A, B**). Starting 1 week after surgery, all mice underwent a battery of behavioral tests to assess the functional effects of hNPC transplantation. After behavior testing, mice were euthanized 5-6 weeks post-transplant surgery to assess engrafted cell placement and fate (**Fig. 2A**). For the initial studies, male R6/2 mice and their age matched WT littermates were assigned to the following groups: 1) WT + vehicle (veh); 2) R6/2 + veh; 3) R6/2 + hNPC; 4) R6/2 + hNPC overexpressing NT-3 (R6/2 hNPC-NT-3); 5) R6/2 + hNPC overexpressing BDNF (R6/2 hNPC-BDNF).

**Figure 2.**
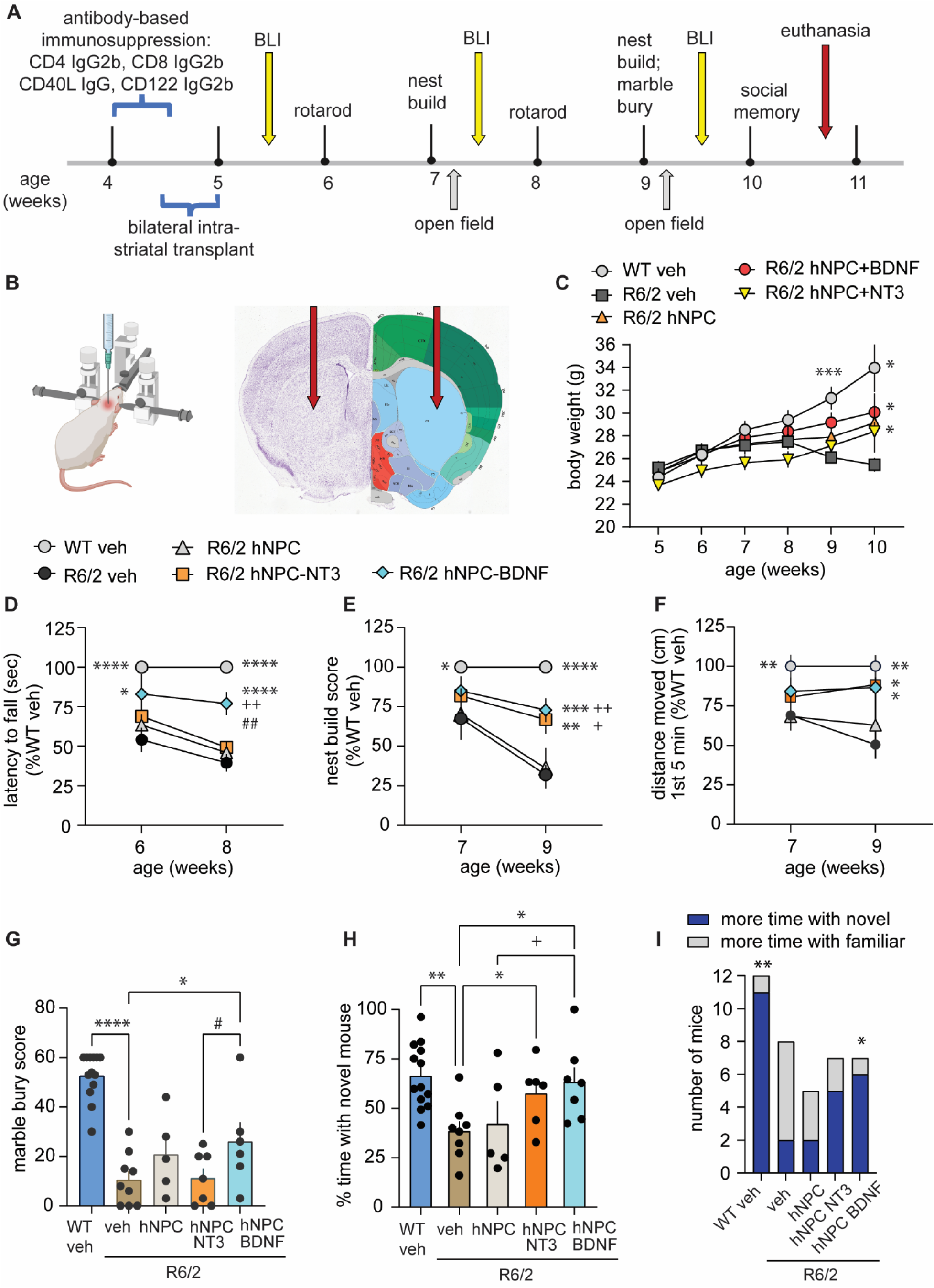
Intrastriatal transplants of hNPCs overexpressing NTs prevent functional deficits in R6/2 mice. (**A**) Experimental timeline of transplantation surgery and behavioral assays in R6/2 mice. An immunosuppression protocol utilizing cocktail of antibodies was administered to all mice via daily intraperitoneal injections starting 2 days before surgery. hNPCs were transplanted bilaterally into the striatum when mice were 4.5 to 5 weeks of age. Bioluminescent imaging (BLI) was performed 1, 3, and 5 weeks post-transplant. Starting 1 week after transplant or sham (veh) surgery, a battery of behavior tests was performed. (**B**) Mouse brain atlas plate showing the targeted site of the transplant. Schematic was created with BioRender.com; Allen Institute mouse brain atlas figure: http://atlas.brain-map.org/atlas. (**C**) Body weights of the WT and R6/2 mice from 5 to 10 weeks of age. Statistical significance was determined with a repeated measures ANOVA and Fisher’s LSD. *p ≤ 0.05 and *** p≤ 0.0005 versus R6/2-veh. (**D**) *Rotarod testing.* At 6 and 8 weeks of age, male R6/2-veh mice fell from an accelerating rotarod faster than WT-veh mice. This deficit was prevented in the R6/2 hNPC-BDNF group, which performed better the R6/2 hNPC and hNPC-NT3 groups 3 weeks post-transplant. (**E**) *Nest building.* At 7 and 9 weeks of age, WT-veh mice had higher nest scores than R6/2-veh mice and, 4 weeks after transplant, R6/2 hNPC-BDNF and R6/2 hNPC-NT3 had higher nest scores than both R6/2 hNPC and R6/2- veh mice. (**F**) *Open field.* At 7 and 9 weeks of age, WT-veh mice traveled further in an open field than R6/2-veh mice and, 4 weeks after transplant, R6/2 hNPC-BDNF and R6/2 hNPC-NT3 mice moved a greater distance than both R6/2 hNPC and R6/2-veh mice. (**G**) *Marble bury.* At 9 weeks of age, WT-veh mice had higher marble bury scores than R6/2-veh mice and R6/2 hNPC-BDNF performed better than both R6/2-veh and R6/2 hNPC-NT-3 mice. (**H, I**) *Social memory.* At 10 weeks of age, R6/2-veh mice spent less time with a novel mouse than WT-veh mice and these deficits were alleviated in R6/2 hNPC -BDNF and -NT-3 mice (**H**). A greater number of WT-veh mice spent more time with the novel mouse than the familiar one while a greater number of R6/2- veh mice spent more time with a familiar than a novel mouse. More R6/2 hNPC-BDNF mice preferred the novel mouse (**I**). Number of mice per group: WT-Veh *n* = 11-13; R6/2-veh *n* = 8-10; R6/2 hNPC *n* = 4-5; R6/2 hNPC-NT-3 *n* = 6-7 and R6/2 hNPC-BDNF *n* = 6-7. Multiple cohorts of mice were used and the data was normalized to the WT-veh group of each cohort for **D-F**. Results are expressed as mean ± s.e.m. All data was normally distributed, as indicated by the Kolmogorov-Smirnov test, and for **D-I**, statistical significance was determined with an ANOVA and Fisher’s LSD. *p ≤ 0.05, ** ≤ 0.01, *** ≤ 0.001 and ****p ≤ 0.0001 versus R6/2-veh; ^+^p ≤ 0.05, ^++^p ≤ 0.005 and ^+++^p ≤ 0.001 versus R6/2 hNPC; and ^#^ p = 0.04 and ^#^ ^#^p = 0.001 versus R6/2 hNPC-NT3.

The body weights of R6/2 mice declined with age compared to WT mice. The intrastriatal implants of hNPCs overexpressing BDNF or NT3 ameliorated this decline by 5 weeks after implant (**Fig. 2C**). Motor balance and coordination was assessed with an accelerating rotarod at 6 and 8 weeks of age (1 and 3 weeks after transplant). R6/2-veh mice fell from the rotarod faster than WT-veh mice at both testing time points (**Fig. 2D**). R6/2 mice transplanted with hNPCs overexpressing NT3 or hNPCs without NT overexpression did not differ in rotarod performance from R6/2-veh mice. However, R6/2 mice transplanted with hNPCs overexpressing BDNF had a longer latency to fall from the rotarod than R6/2-veh mice at 6 and 8 weeks of age and by the last week of testing performed better than mice in the R6/2-hNPC and R6/2-hNPC-NT3 groups (**Fig. 2D**).

Nest construction is a complex, naturalistic, goal-directed behavior involving coordinated manipulation of nesting material and can indicate the motor capabilities and well-being of mice ^30^. At 7 and 9 weeks of age (2 and 4 weeks post-transplant), we assessed nest building by scoring nests based on their height and shape. R6/2-veh mice had lower nest scores than WT-veh mice at both testing times (**Fig. 2E**). At the last test, R6/2 mice with hNPCs overexpressing BDNF or NT3 performed equally well and had higher nest scores than R6/2 mice given vehicle or hNPCs without NT overexpression (**Fig. 2E**).

Motor function was further examined by assessing locomotion in an open field and marble burying. At 7 and 9 weeks of age, R6/2-veh mice travelled less distance in an open field during the first 5 min of a 15 min test than WT-veh mice (**Fig. 2F**). R6/2 mice transplanted with hNPCs overexpressing NT3 or BDNF moved a greater distance than R6/2-veh mice during the last test. At 9 weeks of age, R6/2-veh mice buried fewer marbles and covered them less completely than WT-veh mice (**Fig. 2G**). The R6/2-hNPC-BDNF mice had a higher marble bury score than both the R6/2-veh and -hNPC-NT3 mice. R6/2 mice that received hNPCs without NT overexpression did not show improvements on either of these motor tests.

The striatum plays a role in learning and memory in social contexts so we investigated the effect of intrastriatal hNPC transplantion on social cognition in R6/2 mice ^31^. At 10 weeks of age, R6/2-veh mice spent less time with a novel stimulus mouse than a familiar mouse compared to WT-veh mice suggesting a discrimination deficit as mice tend to prefer social novelty (**Fig. 2H**). R6/2 mice transplanted with hNPCs overexpressing NT3 or BDNF spent more time with the novel versus familiar mouse and, for the R6/2-hNPC-BDNF group, this difference was present between the R6/2 mice given hNPCs without NT overexpression. The latter mice did not differ from R6/2- veh mice. A greater number of WT-veh and R6/2-hNPC-BDNF mice showed a preference for the novel mouse compared to R6/2-veh mice (**Fig. 2I**).

### Transplanted hNPCs survive and proliferate in the striatum of R6/2 mice

The hPSCs used in this study were also engineered to express akaluciferase allowing longitudinal *in vivo* tracking of the survival and proliferation of the transplanted cells via noninvasive bioluminescent imaging. R6/2 mice were imaged 1, 3, and 5 weeks after receiving intrastriatal transplants of hNPCs. The antibody-based immunosuppression paradigm enabled survival and proliferation of each of the transplanted cell types, as evidenced by an increase in bioluminescent signal from week 1 to 5 (**Fig. 3A, B**).

**Figure 3.**
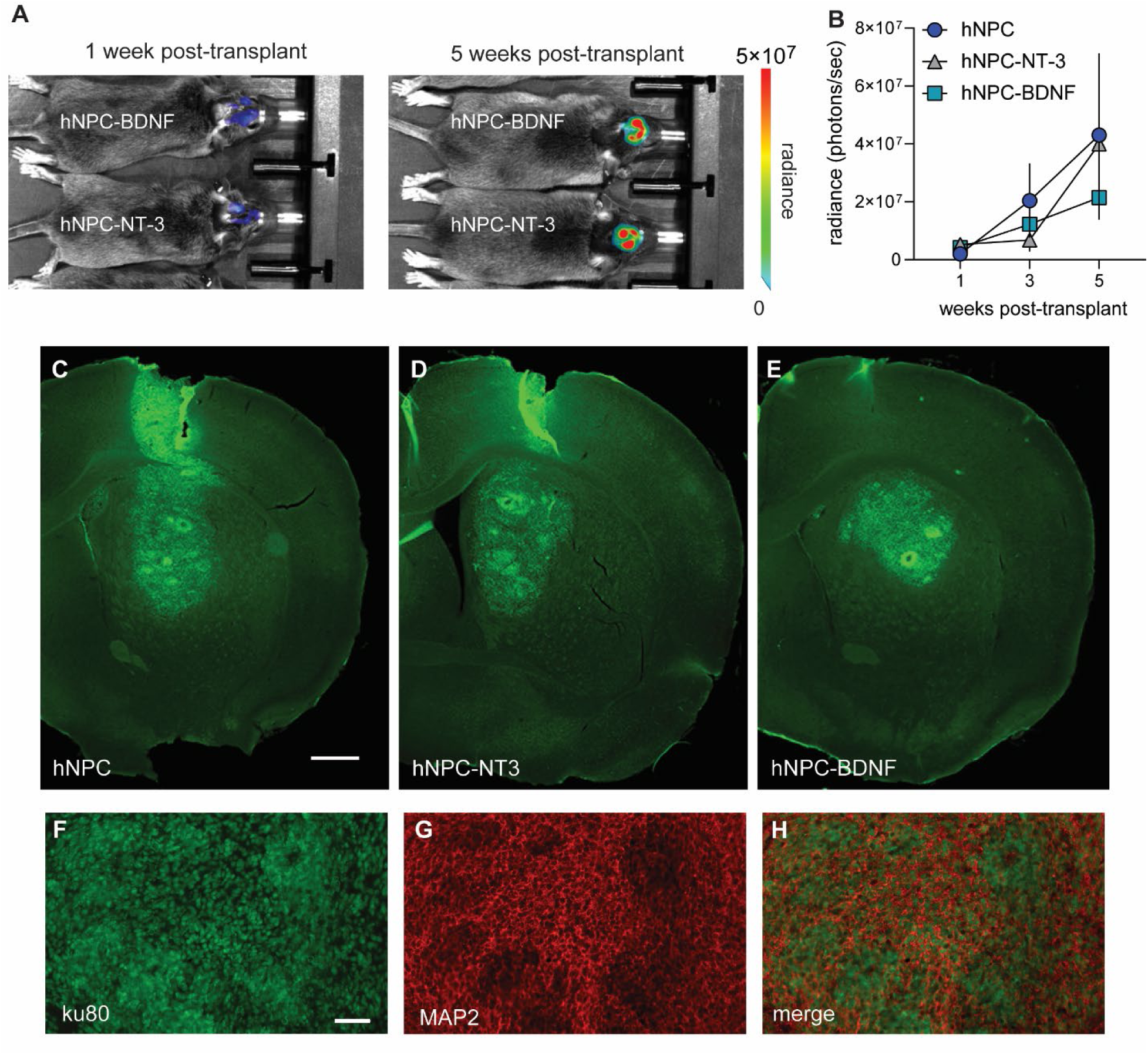
Intrastriatal transplants of hNPCs overexpressing NTs survive and proliferate in R6/2 mice. (**A**) Representative bioluminescent images from one R6/2 hNPC-BDNF mouse and one R6/2 hNPC-NT-3 mouse 1 week after transplant and the same mice 5 weeks after transplant showing that the total emission (radiance) increases from the first to last imaging session. (**B**) Line graph showing the quantification of the total emission during bioluminescent imaging at 1, 3 and 5 weeks after transplant. R6/2 hNPC *n*= 3 mice; R6/2 hNPC-NT-3 *n*= 4 mice, and R6/2 hNPC-BDNF *n*= 5 mice. (**C-E**) Immunostaining for the human nuclei marker, Ku80, indicating engraftment of hNPC (**C**), hNPC-NT3 (**D**), and hNPC-BDNF (**E**) cells in the R6/2 striatum. The engraftment area encompasses a large part of the striatum and contains circular structures resembling rosettes. Scale bar in C = 300 mm and applies to C-E. (**F-H**) Immunostaining for human nuclei Ku80 (**D**) and MAP-2 (**E**) largely overlap (**H**) suggesting the majority of the implanted hNPC-BDNF cells remain neuron-restricted progenitor cells. Scale bar in F = 50 mm and applies to F-H.

WT and R6/2 mice were euthanized 5-6 weeks post-transplant or sham surgery and their brains were collected to assess the location and fate of the injected hNPCs. Immunostaining for the human nuclei marker (Ku80) revealed that engrafted hNPCs were located primarily in the striatum and along the injection needle tract in the cortex. Some hNPCs appeared to migrate along the corpus callosum/external capsule between the striatum and the cortex (**Fig. 3C-E**). Regardless of the cell type, the human cell engraftment site was dense and encompassed a large part of the striatum (**Fig. 3C-E**), which is in accordance with the proliferation evident from the bioluminescent imaging. The engraftment areas of each cell type contained radially arranged cells resembling rosettes indicating that the cells in these formations likely retain their stemness and proliferation potential (**Fig. 3C-E**). Moreover, most of the transplanted cells showed immunostaining for microtubule-associated protein 2 (MAP-2) suggesting they remained neuron-restricted progenitor cells (**Fig. 3F-H**). These results, seen just 5-6 weeks after implant, are consistent with previous studies showing that after transplantation human neural stem cells can remain poorly differentiated for up to 8 weeks ^32,33^.

### Differentiation of BDNF overexpressing hNPCs into striatal progenitors

To test if using a more mature and differentiated neural cell type to deliver NTs would help reduce or prevent the excessive proliferation seen with hNPC engraftment *in vivo*, we further differentiated the hNPCs into striatal progenitors. For this differentiation, we utilized dual SMAD and WNT inhibition to derive hNPCs, as described above, and used a previously described protocol for derivation of striatal progenitors ^34^. hNPCs were treated with recombinant activin A protein, a WNT inhibitor, and a retinoid X receptor (RXR) agonist for 10 days to obtain striatal progenitors (STRpcs) (**Fig. 4A**). Since R6/2 mice transplanted with hNPCs overexpressing BDNF performed better on more of the behavioral assays than R6/2 mice given hNPCs expressing NT3, we derived hSTRpcs from BDNF-C2 cells and mock hPSCs for further transplantation studies. hSTRpcs expressed the striatal markers GSX2 and DLX2 and the forebrain marker FOXG1, as indicated by immunostaining (**Fig. 4B**). Quantification showed that about 80, 55, and 75% of the hSTRpcs were positive for GSX2, DLX2 and FOXG1, respectively (**Fig. 4C**). The mean BDNF concentration in the supernatant from hSTRpcs derived from BDNF-C2 cells was approximately 38 pg/ml per 100K cells, as evidenced by ELISA, and was not detectable in the supernatant from mock cells indicating a lack of BDNF expression (**Fig. 4D**).

**Figure 4.**
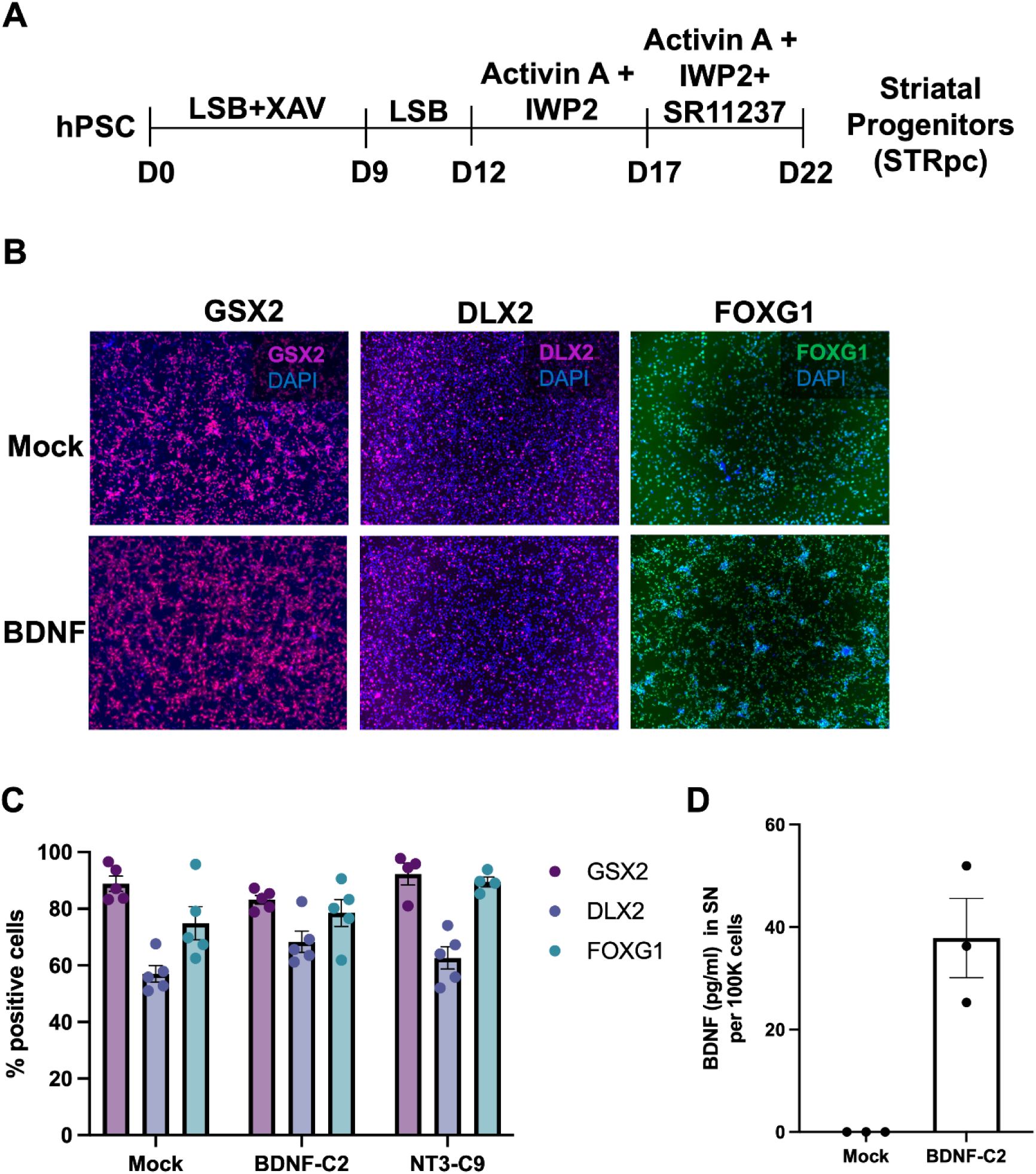
Differentiation of BDNF overexpressing hPSCs into striatal progenitors. (**A**) Schematic of protocol for differentiation of hPSCs into striatal progenitor cells (hSTRpcs). LSB denotes LDN193189 (BMP inhibitor) and SB431542 (TGF-β inhibitor), XAV denotes XAV939 (WNT inhibitor), IWP2 (WNT inhibitor), SR11237 (RXR agonist). (**B**) Representative immunostaining images for striatal progenitor markers GSX2, DLX2 and FOXG1 in hSTRpcs derived from mock and BDNF-C2 ES cell lines. (**C**) Frequency of GSX2, DLX2 and FOXG1 positive hSTRpcs derived from mock and BDNF-C2 hPSCs. Immunostaining images were quantified as area of immunostaining for the corresponding marker relative to the area of DAPI staining and represented as percentage of positive cells. (**D**) Quantification of the ELISA for BDNF in the supernatant (SN) of hSTRpcs derived from BDNF-C2 hPSCs. Mock hSTRpcs were used as a negative control. Data is shown as the concentration of BDNF in SN per 100K cells.

### Intrastriatally transplanted hSTRpcs overexpressing BDNF prevent functional deficits in R6/2 mice

The transplantation studies with hSTRpcs used the same study design as described above (**Fig. 3A**) except that both male and female mice were used. Disrupted motor balance and coordination in R6/2 mice, as assessed with a rotarod, was prevented by intrastriatal transplantation of both hSTRpcs overexpressing BDNF or those without NT overexpression (**Fig. 5A**). Transplanting hSTRpcs overexpressing BDNF also ameliorated deficits in nest building (**Fig. 5B**), ambulation (**Fig. 5C**), goal-directed digging (marble bury; **Fig. 5D**), and rearing (**Fig. 5E**) and grooming (**Fig. 5F**) in a novel environment compared to R6/2-veh mice. hSTRpcs without NT overexpression did not significantly affect any of these behaviors (**Fig. 5**). Moreover, R6/2 hSTRpc-BDNF mice had higher marble bury scores, rearing frequency, and normalized grooming behavior compared to R6/2 hSTRpc mice.

**Figure 5.**
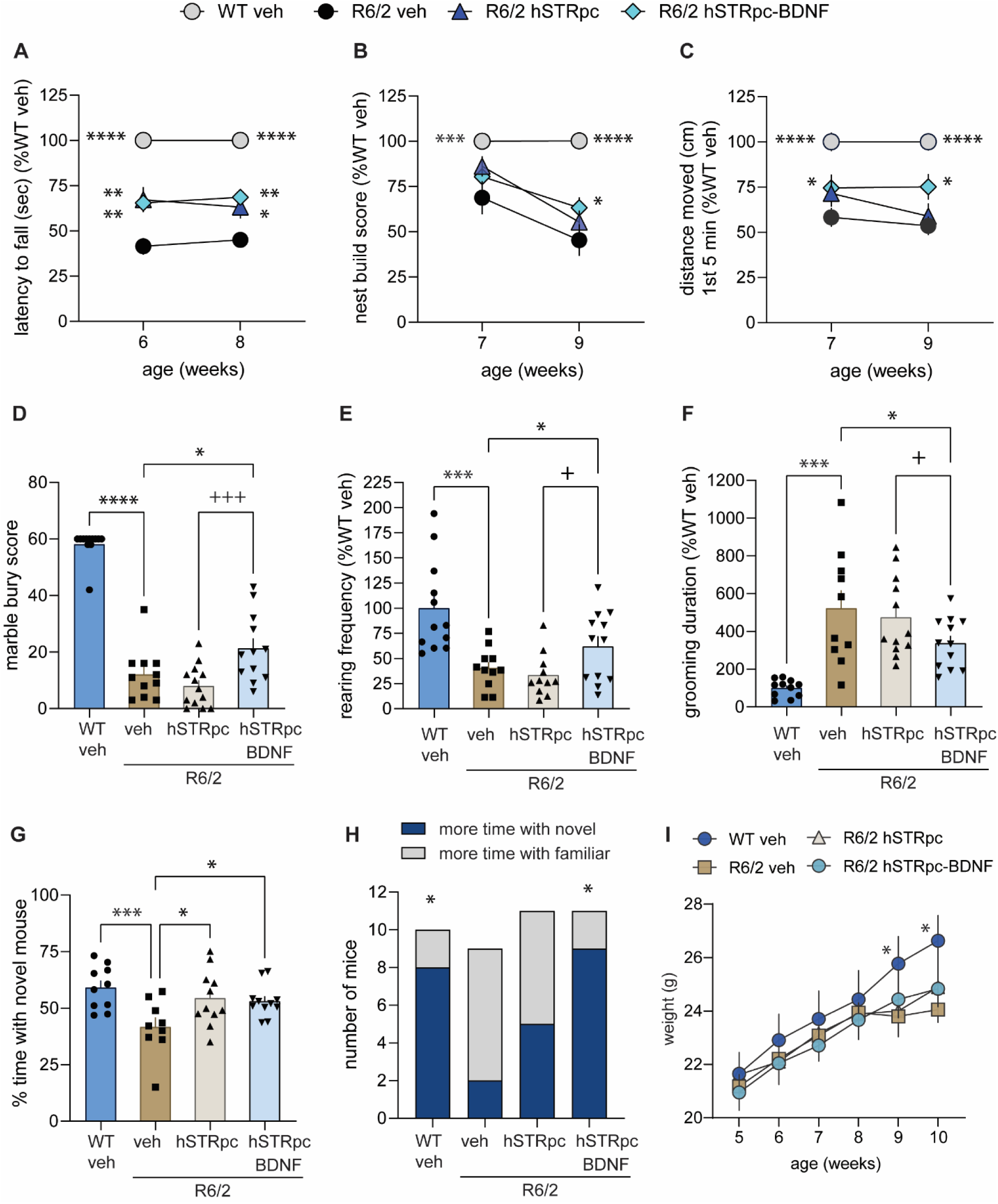
Intrastriatal transplants of hSTRpcs overexpressing BDNF prevent motor and cognitive deficits in male and female R6/2 mice. (**A**) *Rotarod.* R6/2-veh mice had rotarod deficits at 6 and 8 weeks of age. This deficit was prevented in R6/2 mice that received hSTRpc or hSTRpc-BDNF transplants. (**B - F**) Compared to WT-veh mice, R6/2-veh mice showed behavioral disturbances in nest building (**B**), distance traveled in an open field (**C**), marble burying (**D**), rearing (**E**), and grooming (**F**). R6/2 hSTRpc-BDNF mice performed better on each of these behavior tests than R6/2-veh mice (**B-F**) and better than R6/2 hSTRpc on the latter three assays (**D-F**). (**G, H**) *Social memory.* R6/2-veh mice spend less percent time with a novel mouse than WT-veh mice. These social recognition deficits in R6/2 mice were alleviated by intrastriatal transplants of hSTRpc or hSTRpc-BDNF (**G**). A greater number of WT-veh mice spent more time with the novel mouse than the familiar one while a greater number of R6/2-veh mice spent more time with a familiar than a novel mouse. A larger number of R6/2 hSTRpc-BDNF mice preferred the novel mouse (**H**). (**I**) Body weights of the WT and R6/2 mice from 5 to 10 weeks of age. Number of mice per group: WT-Veh *n* = 10-12 mice; R6/2-veh *n* = 9-11; R6/2 hSTRpc *n* = 11-13; R6/2 hSTRpc-BDNF *n* = 12-14. Multiple cohorts of mice were used and the data was normalized to the WT-veh group of each cohort for **A-C,E,F**. Results are expressed as mean ± s.e.m. All data was normally distributed, as indicated by the Kolmogorov-Smirnov test, and statistical significance was determined with an ANOVA and Fisher’s LSD except for the body weights for which a repeated measures ANOVA was used. *p ≤ 0.05, ** ≤ 0.01, *** ≤ 0.001 and ****p ≤ 0.0001 versus R6/2-veh; ^+^p ≤ 0.05 and ^+++^p ≤ 0.001 versus R6/2 hSTRpc.

Social recognition deficits in R6/2 mice were also alleviated by intrastriatal hSTRpc transplantation as indicated by the greater percent time that R6/2 hSTRpc-BDNF and R6/2 hSTRpc mice spent exploring a novel versus familiar mouse compared to R6/2-veh mice (**Fig. 5G**). A greater number of WT-veh mice spent more time with the novel mouse than the familiar one, while R6/2-veh mice did not have a preference for social novelty and a greater number of them spent more time with a familiar than a novel mouse (**Fig. 5H**). Similar to WT-veh mice, a greater number of R6/2 hSTRpc-BDNF mice preferred the novel mouse, whereas this was not the case for R6/2 hSTRpc mice (**Fig. 5H**). The body weights of the R6/2 mice with hSTRpc or hSTRpc-BDNF transplants were not significantly different from R6/2-veh mice (**Fig. 5I**).

### Transplanted hSTRpcs proliferate, differentiate, and produce BDNF

Transplanted hSTRpcs and hSTRpcs overexpressing BDNF were longitudinally tracked *in vivo* with bioluminescent imaging at 1, 3 and 5 weeks after intrastriatal injections. Both cell types survived and proliferated (**Fig. 6A, B**). At 5-6 weeks post-transplant, the brains from mice in all treatment groups were collected so that the location of the human cell engraftment and BDNF levels could be assessed. The engrafted hSTRpcs with or without BDNF overexpression were concentrated around the injection needle tract primarily in the striatum and some were in the corpus callosum and cortex. The engraftment area was smaller than seen with hNPCs and the circular configurations of cells resembling rosettes were not detected in the mice that received hSTRpc or hSTRpc-BDNF transplants (**Fig. 6C-E; Suppl. Fig. 3A-F**). Some transplanted cells migrated along the corpus callosum and distal from the injection rostro-caudally within the striatum.

**Figure 6.**
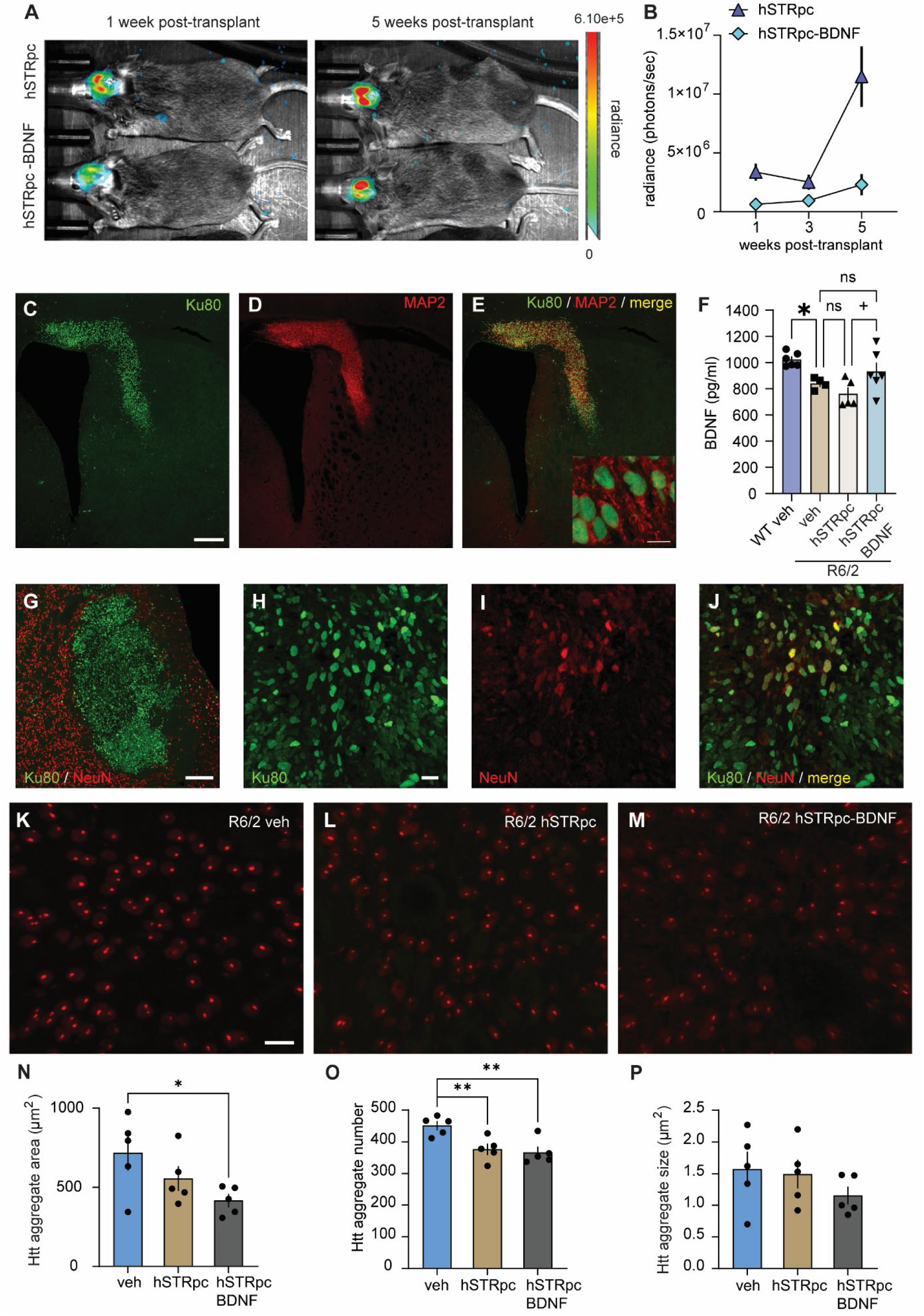
Intrastriatal hSTRpc transplants proliferate, differentiate, produce BDNF, and reduce mHtt aggregates in R6/2 mice. (**A**) Representative bioluminescent images from one R6/2 hSTRpc and one R6/2 hSTRpc-BDNF mouse 1 week after intrastriatal transplant surgery and the same mice 5 weeks post-transplant showing that the implanted cells survive and that the total emission (radiance) increases from the first to last imaging session. (**B**) Line graph showing the quantification of the total emission during bioluminescent imaging at 1, 3 and 5 weeks after transplant. R6/2 hSTRpc *n* = 12 mice; R6/2 hSTRpc-BDNF *n* = 14 mice. (**C-E**) Representative photomicrographs of the STRpc-BDNF engraftment area in the striatum of an R6/2 mouse immunostained for the human nuclei marker, Ku80 (**C**, green), microtubule-associated protein 2 (MAP2) (**D**, red), and the merged channels (**E**, yellow). The engraftment areas were smaller than those seen with hNPCs (see Fig. 4) and the circular cell patterns were not detected in the mice that received hSTRpc (**Suppl. Fig. 3**) or hSTRpc-BDNF transplants (**C-E**). Most of the hSTRpc cells colocalize with the neuron-restricted progenitor marker, MAP2 (**E**, yellow); see also the insert with higher magnification (63X, 300% zoom, Imaris). Scale bar in C = 200 μm and applies to C-E. Scale bar in E (insert) = 5 μm. (**F**) Bar graph showing quantification of mature BDNF levels, as assessed via ELISA, in the striatum of 10-11 week-old male and female mice. Number of mice per group: WT-Veh *n* = 6 mice; R6/2-veh *n* = 4; R6/2 hSTRpc *n* = 5; R6/2 hSTRpc-BDNF *n* = 6. All data was normally distributed, as indicated by the Kolmogorov-Smirnov test, and statistical significance was determined with an ANOVA and Fisher’s LSD. One mouse in the R6/2 hSTRpc group was removed as a statistical outlier (with BDNF levels at 1208 pg/ml). *p = 0.016 WT-veh versus R6/2-veh; ^+^p = 0.0165 R6/2 hSTRpc versus R6/2 hSTRpc-BDNF. ns = not statistically significant. (**G-J**) Immunostaining for human nuclei (green) and NeuN (**G-J**), astrocytes (GFAP; **Suppl. Fig. 3G-J**) or oligodendrocytes (O2;**Suppl. Fig. 3K-N**) showing that few of the transplanted cells differentiate into these lineages. Scale bar in G = 100 μm. Scale bar in H = 10 μm and applies to H -J. (**K-M**) Representative photomicrographs of mHtt immunostaining in the striatum of R6/2 mice with a vehicle injection (**K**) or that were transplanted with hSTRpcs (**L**) or hSTRpcs with BDNF overexpression (**M**). Scale bar in K = 10 μm and applies to K-M. (**N-P**) Bar graphs showing the quantification of the total area (**N**), number (**O**), and size (**P**) of intranuclear mHtt aggregates per a 290 × 290 µm analysis field adjacent to the engraftment area or vehicle injection site (*n*= 5 mice / group). All data was normally distributed, as indicated by the Kolmogorov-Smirnov test, and statistical significance was determined with an ANOVA and Fisher’s LSD. *p = 0.05 and ** ≤ 0.01 versus R6/2-veh group.

Mature BDNF levels were reduced in the striatum of 11-week-old R6/2 mice compared to WT mice given intrastriatal vehicle injections and were not significantly altered in R6/2 mice that received intrastriatal hSTRpc transplants that did not overexpress BDNF (**Fig. 6F**). In contrast, hSTRpc-BDNF transplants increased mature BDNF levels in R6/2 striatum compared to hSTRpc transplants but not vehicle injections. Striatal BDNF levels in R6/2 mice with hSTRpcs overexpressing BDNF did not differ from WT veh mice.

The fate and differentiation profile of the engrafted hSTRpcs was also evaluated using immunostaining for markers for human nuclei (Ku80), the immature neuronal markers doublecortin or MAP2, and the main neural lineages: neurons (NeuN), astrocytes (GFAP), or oligodendrocytes (O4). Most of the engrafted hSTRpcs and hSTRpcs overexpressing BDNF were immunostained with the early neuronal markers, doublecortin (**Suppl. Fig. 3A-C**), or MAP-2 (**Fig. 6C-E, Suppl. Fig. 3D-F**) indicating that most remained neuron-restricted progenitors. A small percentage of the transplanted hSTRpc and hSTRpc-BDNF cells differentiated into post-mitotic neurons (NeuN; 1.95 ± 0.97% of hSTRpc; 2.82 ± 1.41% hSTRpc-BDNF; mean ± s.e.m.) (**Fig. 6G-J**), astrocytes (GFAP; 3.64 ± 1.82% hSTRpc; 4.92 ± 2.46% hSTRpc-BDNF) (**Suppl. Fig. 3G-J**), or oligodendrocytes (O4; 1.52 ± 0.42% hSTRpc; 3.68 ± 1.53% hSTRpc-BDNF) (**Suppl. Fig. 3K-N**). The differentiated cells tended to be on the edges of the engraftment area where the density of the engrafted cells was low.

### Transplanting hSTRpcs with or without BDNF overexpression reduces intranuclear huntingtin aggregates in the R6/2 striatum

A characteristic HD pathology is the intranuclear accumulation of Htt which occurs throughout the brain of HD patients and mouse models. WT Htt is normally present in the cytoplasm of cells but mHtt is abnormally processed and translocates into the nucleus where it aggregates ^28,35^. Htt aggregate load can be reduced by increasing BDNF/TrkB signaling ^3,12,36,37^. We evaluated whether transplanting hSTRpcs that overexpress BDNF into the striatum could reduce the formation of intranuclear Htt aggregates in the striatal area near the engraftment site (**Fig. 6K-M**). R6/2-veh mice have numerous, large intranuclear mHtt aggregates (**Fig. 6K**). The total area and number of aggregates, but not the size, were reduced in R6/2 mice that received intrastriatal transplants of hSTRpcs expressing BDNF (**Fig. 6M, N-P**); the aggregate number was also decreased in the R6/2 mice with hSTRpcs without BDNF overexpression (**Fig. 6L, O**). Intranuclear Htt aggregates were not present in the engrafted cells (**Suppl. Fig. 3O-Q**).

### Activating the orthogonal safeguard eliminates engrafted hSTRpcs overexpressing BDNF

To validate the application of the safety switch expressed in the gene-targeted hPSCs, dimerizers such as AP21967 or rapamycin can be used to activate caspase 9 dimer formation and trigger apoptosis of cells. We tested whether AP21967 or rapamycin could activate the ACTB safety switch *in vitro* to eliminate all cell types. Both AP21967 and rapamycin (1 nM) almost completely eliminated the hNPCs derived from mock and BDNF-C2 hPSCs (**Supp Fig. 4A**). Rapamycin at up to 8-fold lower concentrations (0.125 nM) also activated the safety switch and effectively eliminated the hSTRpc-BDNF cells (**Supp Fig. 4B**).

Next, we investigated the efficacy of the safety switch *in vivo* after transplanting the orthogonal safeguard gene-edited hSTRpcs that overexpress BDNF into the striatum of R6/2 mice. Ten days after surgery, bioluminescent imaging was performed to establish hSTRpc-BDNF engraftment (**Fig. 7A**). We used rapamycin to activate the safety switch as it crosses the blood brain barrier more effectively than AP21967^38^. Rapamycin (20 mg/kg) or vehicle was administered via i.p injection daily for 3 days, and imaging was performed again 2 days later to assess safety switch activation. Rapamycin effectively eliminated the bioluminescent signal, while the signal was still strong in R6/2 mice given vehicle (**Fig. 7A, B**). Moreover, 7 days post-injection, the hSTRpc-BDNF cells survived in R6/2 mice that received vehicle as Ku80 staining for human nuclei was prevalent and there was little or no staining for Fluoro-JadeB (FJB), which stains degenerating neurons (**Fig. 7C, D**). In contrast, the R6/2 hSTRpc-BDNF mice given rapamycin had FJB staining near the engraftment area and had minimal immunostaining for human nuclei (**Fig. 7E, F**), consistent with the lack of bioluminescent signal post-injection (**Fig. 7A, B**), indicating the death of the implanted hSTRpcs. Many of the degenerated cells may have already been removed 10 days after the rapamycin injections as FJB staining has been shown to be substantially reduced 7 days after an insult ^39^. Rapamycin injection and death of the engrafted cells did not appear to cause an excessive microglial response as indicated by immunostaining for the microglial marker, IBA-1 (**Fig. 7F**). Thus, activating the safety switch with rapamycin may be an effective way to rectify potential unwanted or harmful effects of hPSC transplantations, if they occur.

**Figure 7.**
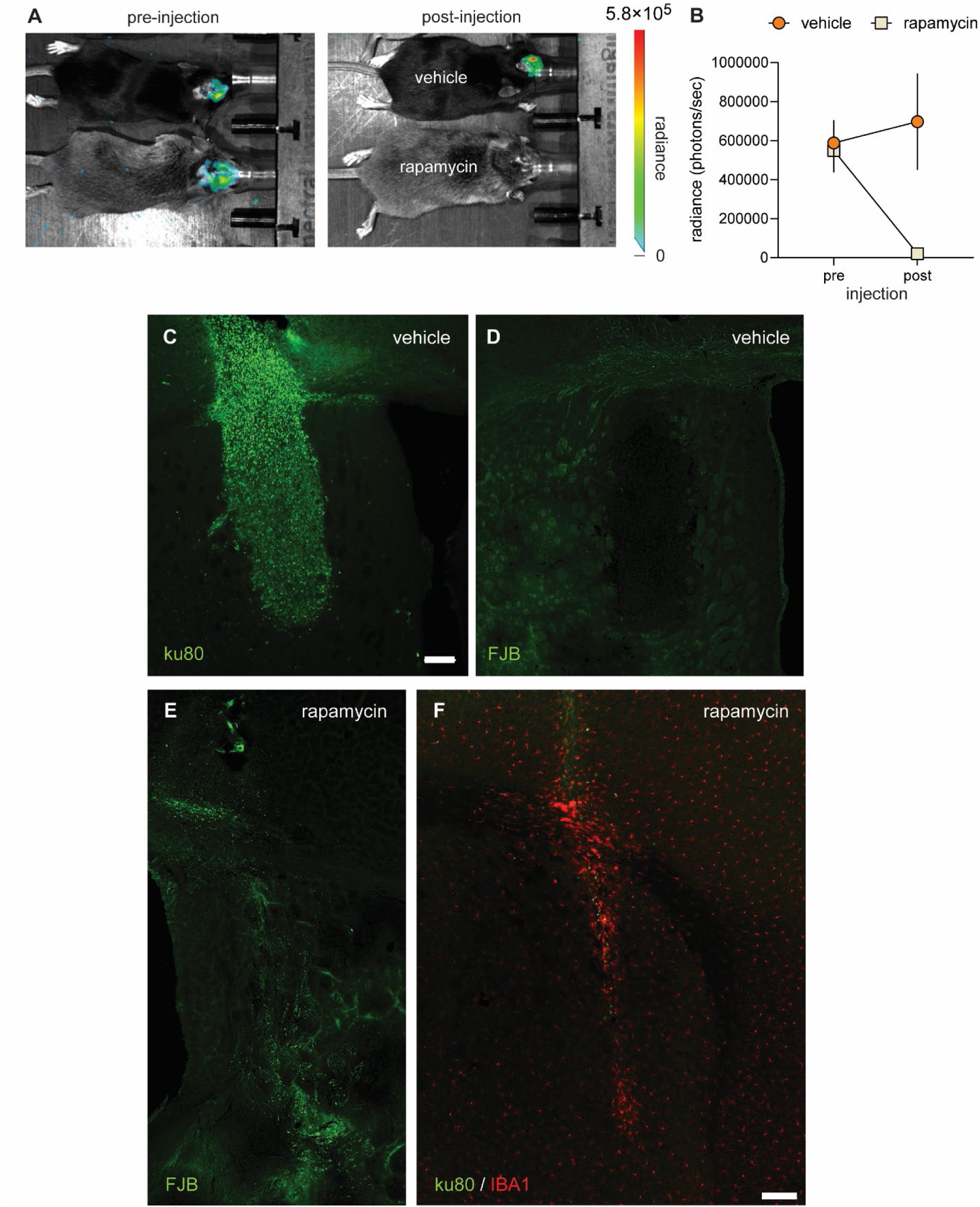
Activation of the safety switch eliminates transplanted hSTR-BDNF progenitor cells. (**A**) Representative bioluminescent images from one R6/2 hSTRpc-BDNF mouse that received vehicle and one that received rapamycin for 3 days starting 10 days after intrastriatal transplant surgery. The left panel is before their vehicle or rapamycin injection and the right panel is the same mice 2 days post-injection. (**B**) Line graph showing the quantification of the total emission before and after vehicle (*n*= 2 mice) or rapamycin injection (*n*= 7 mice) in R6/2 mice. (**C, D**) Immunostaining for the human nuclei marker, Ku80, (**C**) and FluoroJadeB (FJB; **D**) in the striatum showing that the transplanted cells survived in a representative R6/2 hSTRpc-BDNF mouse that received vehicle. (**E, F**) FJB staining near the striatal injection site of R6/2 hSTRpc-BDNF mice given rapamycin (**E**) and the minimal immunostaining for ku80 (green) (**F**) is consistent with death of the implanted hSTRpcs. Some immunostaining for the microglial marker, IBA-1 (red), aggregated near the injection tract (**F**). Scale bars = 100 mm.

## Discussion

The effects of stem-cell based strategies for neuroregeneration and NT delivery have been assessed in many HD preclinical studies as well as in human HD trials ^13,15,40,41^. Initial clinical studies involved transplanting fetal neural stem cells in the caudate-putamen of small cohorts of HD patients ^41^. These trials met with some success in that minimal side-effects regarding surgery or immunosuppression ensued, the grafts largely survived, and, although inconsistent, motor improvement was noted. However, little connectivity between the graft and host tissue occurred and the tissue source raised ethical concerns ^41^. Three recent clinical trials are ongoing each evaluating the efficacy and safety of dental pulp-derived MSCs (Cellavita HD) in HD patients ^41,42^. These cells have been shown to produce BDNF in a chemically-induced rat model of HD and have immunomodulatory properties ^43^. While positive treatment effects were also seen with dental pulp-derived MSCs, these cells have limited proliferation potential and poor differentiation ability. They carry the safety risks of non-directional differentiation and tumor formation ^42^. Thus, stem-cell therapy remains a very promising treatment strategy for HD especially given the regional specificity of neuronal loss however its practicality, reliability, and safety have yet to be established in human trials.

hPSC-based therapies are currently being tested in preclinical studies for neurodegenerative diseases since hPSCs have the potential for derivation of a nearly unlimited amount of therapeutically relevant neural cell types ^44^. hPSC-derived neural stem cells (NSCs) are being tested in an ongoing clinical trial for Parkinson’s disease following extensive preclinical studies showing therapeutic efficacy in animal models of the disease ^15,45^. Few other preclinical studies have investigated the effectiveness of hPSCs for HD ^40^. Most of these studies relied on the differentiation of the transplanted hPSC-derived NSCs into neurons *in vivo* to provide neurotrophic support and potentially replace degenerating neurons. The studies offer potential as regenerative therapeutic strategies as they show that the engrafted cells formed synaptic connections, improved synaptic plasticity, and ameliorated mHtt-related transcriptional deficits in HD mouse striatum to ameliorate behavioral dysfunction and neuropathology ^13,16,18,19^. The current study used hPSCs already differentiated into hSTRpcs overexpressing BDNF and showed that their intrastriatal transplantation reduced HD-related functional deficits and mHtt aggregate formation in R6/2 mice. Although post-mitotic neurons were sparse amongst engrafted BDNF-hSTRpcs, some appeared to be differentiating along a neuronal lineage. Thus, providing more stable and constitutive NT support through transplantation of neural cells overexpressing BDNF could further improve the efficacy of a hPSC-based therapy for HD. Another novel finding of this study was that using orthogonal safeguards can remove a major safety risk associated with PSC-based therapies in that, if need be, engrafted cells can be eliminated after intracranial transplantation, without signs of reactive gliosis, which would be essential in the case of excess proliferation or any other potential hazards associated with hPSC-based therapies. These results provide further support for hPSC-based neuroprotective treatments for HD and other neurodegenerative disorders.

BDNF delivery may be an important contributor to the prevention of the functional deficits and the reduction of intranuclear aggregates in the R6/2 mice in this study. mHtt depletes BDNF gene expression and disrupts its anterograde transport from the cortex to the striatum causing decreased striatal protein levels of the NT in HD patients and mouse models ^4,5,46^. These BDNF deficits contribute to HD-associated motor and cognitive impairments and neurodegeneration ^3,4^. Administering exogenous BDNF into the striatum or crossing mice that have the HD mutation with BDNF over-expressing mice reduced behavioral deficits and HD pathology, including intranuclear Htt aggregates ^3,4,36,47,48^. These studies indicate that BDNF deficits may play a causal role in the development of characteristic HD pathologies and that restoring the NT can prevent neurodegeneration. Accordingly, elevating striatal BDNF and its signaling in HD mouse models via transgenic methods or with small molecules slows disease progression ^3,4,12^. Moreover, other HD preclinical studies assessing PSC-based therapies have largely attributed the obtained beneficial effects to BDNF produced by the cells and are corroborated by the results we show here ^13,16,18,19^. NT3 may also play a role in preventing HD-related pathogenesis as its levels are reduced in HD patients and mouse models and up-regulating it or activating its signaling provides neuroprotection ^3,11,12^. This initial part of this study showed that R6/2 mice transplanted with hNPCs overexpressing NT3 did alleviate some of the motor deficits compared to those with vehicle, however the effects of hNPCs overexpressing BDNF were more robust. Thus, while it was beyond the scope of this study to conduct a rigorous assessment of both hNPC cell types overexpressing NTs, the positive results seen with hNPCs overexpressing NT3 may suggest further studies with these cells or derivatives that produce more NT3 may be warranted.

Intracranial transplantation of stem cells that produce NTs may circumvent some of the technical challenges faced with attempts at direct CNS administration of NTs, and the bioavailability obstacles that occur with strategies of peripheral administration such as low plasma stability with brief serum half-lives, the need for repeated administration, and limited blood brain barrier permeability. Moreover, PSCs producing NTs may offer a more sustained and targeted delivery than NT infusion with more controlled dosing and fewer side effects. A concern with using hPSCs that constitutively express NTs could be excess NT production leading to detrimental side effects, however the inducible safeguards employed here would be able to resolve this potential setback. Thus, the results presented here support an hPSC-based approach to delivering NTs and offer a viable precaution against safety risks. Additional preclinical validation of this strategy is warranted including evaluating the effects of BDNF-hSTRpcs in a full-length HD mouse model with a slower progressing disease course than R6/2 mice. This slower progression would allow a longer post-transplantation examination period during which more of the engrafted cells could mature into post-mitotic neurons. Although the main objective of this study was cellular NT delivery, it would be of interest to evaluate the potential of these hSTRpcs as a cell replacement therapy which would require evidence that the engrafted cells can establish synaptic connections with the host’s local circuitry. To overcome a possible host immune response against transplanted allogeneic cells, it would be ideal to deliver the NTs through a gene-corrected HD patient-specific iPSC-based platform.

In all, the treatment strategy described here constitutes a next generation for PSC therapies as it goes beyond prior methods against neurodegeneration by integrating multi-plex genome editing to engineer combined features of both efficacy and safety into the therapeutic cell product. Given that HD can be diagnosed before clinical onset and that the neuronal loss is largely region specific and significantly affected by NT loss, hPSC-mediated NT delivery could be a very promising therapeutic strategy. This study provides proof-of-concept for developing a hPSC-based platform for NT delivery as a disease-modifying approach for the treatment of HD and other neurodegenerative diseases with a built-in safeguard against excessive growth or deleterious biological effects of engrafted cells.

## Methods

### Human pluripotent stem cell culture

For this study, we used a previously generated human embryonic stem cell (hESC) line H9 that expresses dual safety switches under the control of NANOG and ACTB promoters. This cell line was also engineered to express the akaluciferase transgene which permits *in vivo* bioluminescent cell tracking ^25^. ESCs were maintained in mTeSR1 medium (STEMCELL technologies, 85850) on Matrigel (Corning, 354277) coated plates. hESCs were passaged once they reach a confluence of 70 to 80% following dissociation with Accutase (Innovative Cell Technologies, AT104) or ReLeSR (STEMCELL technologies, 100-0484). ESCs passaged with Accutase were maintained for 24 hours with mTeSR1 supplemented with 10 μM of the ROCK kinase inhibitor, Y-27632 (Cayman Chemical, 10005583); cells were switched to mTeSR1 medium without Y-27632 after 24 hours. For freezing, hESCs dissociated with Accutase or ReLeSR were pelleted and resuspended in STEM-CELLBANKER (Amsbio, 11924) freezing medium at a density of 250 to 500K cells per 100 μL.

### Genome editing of hPSCs

hPSCs were gene targeted at the HBB safe harbor locus for NT expression using Cas9 RNP and AAV6 based gene editing platform ^49^. Gene editing used a Hifi Cas9 protein (IDT, 1081061 or Aldevron, 9214)^50^ and an sgRNA chemically modified to include 2′-O-methyl-3′-phosphorothioate at the first and last three nucleotides ^51^. The genomic target sequence of the *HBB* sgRNA used was 5’ CTTGCCCCACAGGGCAGTAA 3’. For generation of the AAV6 HDR donor vector, homology arms flanking the gRNA target site and the insert sequence consisting of Ubiquitin C promoter, BDNF or NT3 cDNA and bGH polyA were cloned into pAAV-MCS2 plasmid digested with NotI. The generated transfer plasmid was sequence confirmed and a maxiprep was prepared using PureLink™ Expi Endotoxin-Free Maxi Plasmid Purification kit (Thermo Fisher Scientific, A31231). For AAV6 production, the transfer and pDGM6 packaging plasmids were co-transfected in 293T cells (ATCC) using PEI reagent (Polysciences, 23966-1), as described previously ^52^. 72 hours post-transfection, AAV6 was extracted and purified from 293T using AAVpro purification kit (Takara, 6666) following the manufacturer’s instructions. AAV6 titer was determined using ddPCR, as described previously ^52,53^.

For gene editing, hPSCs were pretreated with mTeSR1 supplemented with 10 μM Y-27632 for 24 hours, and, the next day, hPSCs at a confluency of around 70% were dissociated into single cells with Accutase. The Cas9 and sgRNA RNP complex was formed by combining 5 μg of Hifi Cas9 protein and 1.75 μg of sgRNA and incubating at room temperature for 15 mins. P3 primary cell nucleofector kit (Lonza, V4XP-3032) and 4D nucleofector (Lonza) were used for delivery of RNP into hPSCs. 100K to 500K of the dissociated hPSCs were resuspended in 20 μL of P3 nucleofection solution consisting of the RNP complex and hPSCs were nucleofected using the CA137 program. After nucleofection, hPSCs were plated at a density of 100K cells per well of a 48-well plate in mTeSR1 medium supplemented with 10 μM Y-27632 and AAV6 donor was added at a multiplicity of infection (MOI) of 5K or 10K. On the following day, the cells were switched to fresh mTeSR1 medium with 10 μM Y-27632 and then from the next day, cells were maintained in mTeSR1 medium without Y-27632. To determine the frequency of gene targeting, ddPCR was performed on the genomic DNA. At 4-5 days post gene editing, genomic DNA was extracted from hPSCs using Quick Extract DNA solution (Lucigen, QE09050). Gene targeting specific amplicon was amplified using one primer on the bGH poly A sequence of the insert and the other primer outside the right homology arm. The following are the ddPCR primer/probe sequences used: *HBB* (In-out PCR) (gene targeting amplicon),

Forward Primer (FP): 5’-GGGAAGACAATAGCAGGCAT-3’,

Reverse Primer (RP): 5’-CGATCCTGAGACTTCCACAC-3’,

Probe: 5’-6FAM/TGGGGATGC/ZEN/GGTGGGCTCTATGGC/3IABkFQ-3’

*CCRL2* (reference amplicon),

FP: 5’-GCTGTATGAATCCAGGTCC-3’, RP: 5’-CCTCCTGGCTGAGAAAAAG -3’

Probe: 5’-HEX/TGTTTCCTC/ZEN/CAGGATAAGGCAGCTGT/3IABkFQ -3’

The frequency of gene targeting was measured by normalizing the copies of gene targeting amplicon relative to that of the reference amplicon.

### Single cell cloning of hPSCs and clonal genotyping

To isolate single cell clones, gene targeted hPSCs were plated at a density of 250 cells per well of a 6-well plate in mTeSR1 medium supplemented with 1X CloneR (STEMCELL technologies, 05888) and incubated for 48 hours. On days 3 and 4, cells were switched to fresh mTeSR1 supplemented with 1X CloneR. From day 5, the cells were maintained in mTeSR1 medium without CloneR. At D7-10, single cell hPSC colonies were scraped and propagated individually. To assess the genotype of the single cell hPSC colonies, PCR to amplify the region spanning the knock-in was used. For this, we designed a forward primer annealing upstream of the left homology arm and a reverse primer annealing downstream of the right homology arm. Based on the PCR results, we identified clones with bi-allelic gene targeting. The following are the sequences of primers used for genotyping: FP: 5’-TAGATGTCCCCAGTTAACCTCCTAT-3’; RP: 5’-TTATTAGGCAGAATCCAGATGCTCA-3’.

### ELISA for BDNF and NT3 in cell culture supernatants

To assess the secreted levels of BDNF or NT3, cell culture supernatants were collected from hPSCs, hNPCs, and hSTRpcs and analyzed by ELISA using a human BDNF ELISA Kit (Abcam, ab212166) or a human NT3 ELISA kit (Thermo Fisher Scientific, EHNTF3). Both ELISAs were performed as per the manufacturer’s instructions. A SpectraMax M3 plate reader (Molecular devices) was used to measure the absorbance of the ELISA samples at 450 nm. The concentration of BDNF or NT3 in the supernatant was determined using a standard curve.

### Generation of human neural and striatal progenitor cells

hPSCs were differentiated into NPCs using a previously described protocol ^27^. hPSCs were plated on Matrigel-coated plates at a density of 20K cells per well of a 12-well plate in mTeSR1 medium supplemented with 10 μM Y-27632. The following day, cells were switched to Essential 6 medium (Thermo Fisher Scientific) supplemented with 0.5 μM of LDN193189 (BMP inhibitor, Cayman Chemical-11802), 10 μM of SB431542 (TGF-β inhibitor, Cayman Chemical-13031) and 5 μM of XAV939 (WNT inhibitor, Cayman Chemical-13951) and were cultured in this medium for 9 days with medium changes every other day. At D9, cells were switched to Essential 6 medium supplemented with 0.5 μM of LDN193189 and 10 μM of SB431542 for 3 days to derive neural progenitors.

For differentiation into striatal progenitors (STRpcs), NPCs were plated on 0.001875% polyethylenimine (Sigma, 03880) or Matrigel-coated plates. NPCs were cultured for 5 days in Neurobasal A medium (Thermo Fisher Scientific, 10888022) supplemented with B27 without Vitamin A (Thermo Fisher Scientific, 12587010), 2.5 μM of IWP2 (Cayman Chemical, 13951) and 50 ng/ml of recombinant Activin-A (Peprotech, 120-14P) with media change every other day. Next, cells were cultured in Neurobasal A medium supplemented with B27 without Vitamin A, 2.5 μM of IWP2, 50 ng/ml of recombinant Activin-A and 100 nM of SR11237 (Tocris, 3411) for 5 days with media change every other day to derive STRpcs^34^. For transplantation, neural and striatal progenitors were dissociated with Accutase and resuspended at a density of 100K cells in 2 μL of HBSS.

### Immunostaining of *in vitro* cultured cells and quantification

For immunostaining analysis, cells were fixed with 4% paraformaldehyde (Electron Microscopy Sciences, 15710) in PBS for 20 mins at RT, followed by permeabilization with 0.3% TritonX-100 (Sigma, T8787) in PBS for 20 mins at RT. Cells were then incubated in a blocking solution of 3% BSA in PBS for 1 hr at RT. After blocking, cells were incubated overnight at 4°C in primary antibody diluted in 3% BSA. Following day, cells were washed once with PBS and were incubated in DAPI (Santa Cruz Biotechnology, sc-3598, 1:5000), and secondary antibody diluted in 3% BSA at RT for 45 mins. After a wash with PBS, cells were maintained in fresh PBS and imaged under BZ-X710 microscope (Keyence). Following are the primary antibodies used, anti-OCT3/4 (sc-5279, 1:200), anti-SOX2 (sc-365823, 1:200), anti-NANOG (sc-293121, 1:200) (all from Santa Cruz Biotechnology), anti-PAX6 (DSHB, PAX6-s, 1:50), anti-SOX1 (BD biosciences, BDB562224, 1:100), anti-FOXG1 (Abcam, ab196868, 1:750), anti-GSX2 (Millipore, ABN162,1:200) and anti-DLX2 (SCBT, sc-393879,1:200). Following are the secondary antibodies used: Alexa Fluor 647 goat anti-mouse (Thermo Fisher Scientific, A21235, 1:500) and Alexa Fluor 488 goat anti-rabbit (Thermo Fisher Scientific, A11008, 1:500).

For quantification of immunostaining images from in vitro cultured cells, we used FIJI software. The area of staining for the corresponding markers and DAPI were quantified after manual thresholding and the relative frequency was calculated to determine the percentage of cells positive for the corresponding marker.

### *In vitro* safety switch validation and cell viability assay

To validate safety switch activation in vitro, NPCs and STRpcs were treated with either AP21967 (1 nM) (Takara, 635055) or Rapamycin (1 nM, 0.5 nM, 0.25 nM and 0.125 nM) (Cayman Chemical, 13346) for 2 days. WT hPSC was used as negative control. For assessing cell viability, MTT assay was performed on the cells post AP21967 or Rapamycin treatment. Cells were incubated in 0.5 mg/ml of MTT resuspended in growth medium for 2 hours. Following MTT incubation, cells were lysed using a lysis buffer consisting of 0.1 N HCL, 0.5% SDS in isopropanol. A SpectraMax M3 plate reader (Molecular devices) was used to measure the absorbance at 570 nm with 650 nm as the reference. The absorbance was used to calculate cell viability as percentage relative to the untreated cells.

### *In vivo* study design

This study was designed to determine whether intrastriatal transplantation of NT-overexpressing human pluripotent stem cells (hPSCs) engineered with orthogonal safeguards can provide neuroprotection in an HD mouse model and if activating the safety switches could eliminate the transplanted cells without unwanted degenerative effects. As a first step, we examined if transplanting hPSC-derived neural progenitor cells (hNPCs) that overexpress BDNF or NT-3 into the striatum of R6/2 mice would prevent HD-related behavioral deficits, and if so, which cell type (BDNF or NT-3 overexpressing) would produce the most robust effects compared to hNPCs without NT overexpression. Male R6/2 mice and their male wild-type (WT) littermates were used (*n* = 5-12 mice/group). After identifying the most effective hNPCs, we transplanted further matured striatal neuron progenitors (hSTRpcs) and assessed their efficacy in a follow-up, statistically-powered preclinical efficacy study with more endpoints to evaluate HD phenotypes. This part of the study used both male and female R6/2 and WT mice that were semi-randomly assigned to body-weight balanced groups (*n* = 5-7 male and 4-7 female mice/group). The number of mice contributing to each analysis is provided in the figure captions. A G*power analysis revealed that 10-12 or 4-5 mice/group should be sufficient to obtain statistical significance on behavior and histological assays, respectively, when we anticipate a 15-20% difference between groups and greater variability in the transgenic and experimental groups based on previous R6/2 studies.

### Mice, Husbandry, and Genotyping

All animal procedures were conducted in accordance with the National Institutes of Health Guide for the Care and Use of Laboratory Animals using protocols approved by the Institutional Animal Care and Use Committee at Stanford University. These protocols included efforts to minimize animal suffering and the numbers used. Furthermore, the study protocols were approved by the Stem Cell Research Oversight committee at Stanford University.

Breeding pairs of R6/2 mice were purchased from Jackson Laboratories [female hemizygous ovarian transplant B6CBA-TgN (HD exon1)62; JAX stock #006494]. Males and females from litters born to these breeding pairs (R6/2 mice and WT littermates) were used in this study. Mice were group-housed in a pathogen-free animal facility with a 12-h light-dark cycle (on 6 am, off 6 pm) until after surgery when they were singly housed to prevent cagemates from re-opening surgical incisions. All mice received Enviro-Dri paper strands for nesting material with water and food freely available. Tail DNA was used for genotyping via real-time PCR by TransnetYX Inc. (Cordova, TN) and CAG repeat number measurement via ABI GeneMapper 4.0 by Laragen Inc. (Culver City, CA). R6/2 mice in this study had an average of 123 ± 3.1 (mean ± SD) CAG repeats.

### Immunosuppression protocol

The use of human cells in immunocompetent mice, as is present in the R6/2 mice used here, necessitates an immunosuppression regimen to enable engraftment of the transplanted cells. Here, we used an antibody-based conditioning protocol (modified from ^54^) for immune suppression which targets T cells and natural killer cells. Starting two days (Day -2) before transplantation surgery (Day 0), mice were given a daily intraperitoneal injection of an antibody cocktail consisting of: 100 µg of anti-CD4 IgG2b (clone GK1.5) and 100 µg of anti-CD8 IgG2b (clone YTS169.4), which are T cell surface molecules that mediate T cell recognition and activation. For Day -2, the cocktail also contained 250 µg of CD122 IgG2b (clone Tm-β1), to deplete natural killer cells, and, on Day 0, it also contained 500 µg of anti-CD40L IgG (clone MR-1), which is expressed by activated T cells. All antibodies were InVivo monoclonal Abs purchased from BioXCell (Lebanon, NH).

### Transplantation surgery

Bilateral intrastriatal injections of hNPCs, hSTRpcs, or vehicle [Hank’s balanced salt solution (HBSS)] were performed using a digitized small animal ultra-precise stereotaxic instrument (David Kopf Instruments) and an adjustable stage platform. Mice, aged 4.5 to 5 weeks, were anesthetized with isoflurane (3% induction), injected subcutaneously with a sustained release (48 hr) analgesic [buprenorphine SR (0.5 mg/kg)], and the scalp was cleaned of fur before the mouse was placed in the stereotaxic apparatus. Throughout the surgery, mice were anesthetized with isoflurane maintained at 1-2% in 100% oxygen (0.8 L/min) delivered via a nose cone, their eyes were covered with sterile lubricant to avoid drying, and their body temperature was monitored using a rectal probe thermometer (Physitemp) and maintained (36-37°C) using an electronically-controlled heating pad. Lidocaine (2%) was applied to the scalp before it was incised to expose the skull. Holes were drilled in the skull bilaterally for injection at the following coordinates relative to bregma (mm): anteroposterior, +0.75, mediolateral, ±1.7, and dorsoventral, −3.25. Before each injection, the patency of the needle was checked by dispensing 0.2 μl of solution. Mice were bilaterally injected with either 100,000 hNPCs or hSTRpcs per brain hemisphere (2 μl/injection) or vehicle (2 μl HBSS) using a 5 μl Hamilton syringe with a 33-gauge needle (1/2" length) fixed in a stereotaxic injector (Stoelting QSI) set to inject at 0.5 μl/min. The injection needle remained in place for 3 min after each injection to allow solution diffusion. Next, bone wax was used to cover the skull holes and the scalp incision was sealed with dermabond. Mice were singly housed after surgery and allowed to recover from anesthesia in cages with heating pads.

### Behavior testing

For all behavior tests, mice were handled by the experimenter at least twice before testing and they were habituated to the testing room, which had even and dim lighting, for 30 min prior to testing. Each testing apparatus was thoroughly cleaned between mice with 70% ethanol. Behavior analyses were conducted by experimenters that were blind to treatment and genotype.

#### Rotarod

The motor balance and coordination of R6/2 mice was assessed with a rotarod for 2 consecutive days at 6 and 8 weeks of age. Mice were trained on the first day and received 3 trials at a fixed speed of 15 rpm with a 60 sec maximum duration and an inter-trial interval of 5 min. Mice were tested on the second day and received 2 trials using an accelerating speed (4 to 40 rpm over 300 sec) with a 300 sec maximum duration and an intertrial interval of 5 min. The latency to fall from the rotarod was recorded and the 2 test trials were averaged for each mouse.

#### Nest building

Nest-building was assessed at 6 and 9 weeks of age. Singly housed mice were given 6g of uncondensed nesting material (Enviro-dri®) in the late afternoon and allowed to manipulate the material overnight during the dark phase of the light cycle. The next morning, during the light phase, nests are scored from 1 to 5 based on the nesting material length, width, and height as well as position in the cage relative to the starting point ^30^. A score of 0 is assigned to untouched nesting material and a score of 1 is given if there is no clear nest site (*e.g.* scattered nesting material). A point is designated for each increasingly defined nest to reach a maximum score of 5.

#### Open field

Mice were allowed to freely explore an opaque plexiglass chamber (45 cm L X 45 cm W X 35 cm H) for 15 min. Their behavior was recorded by an overhead camera and then analyzed with Ethovision XT software v15 (Noldus). Distance moved and duration of time in the periphery were measured.

#### Marble bury

The marble bury test was conducted at 9 weeks of age and evaluates digging and exploratory behavior. Twelve marbles (15 mm diameter, evenly spaced in a 4x3 layout) are placed on top of large flake (8/20 size) ASPEN wood chip bedding (5 cm deep) in a standard rat cage (13 ¼ cm L × 10 ½ cm W × 7 ¼ cm H) which is larger than a mouse home cage. Their behavior was recorded for 30 min by an overhead camera (Thorlabs) and analyzed with Ethovision XT software v15 (Noldus). The number and duration of digging events was measured and the number of marbles buried partially (at least ⅓) or fully at some point during the test was recorded. A marble that was 1/3 buried was scored a 1, ½ buried was scored a 3, and fully buried was scored a 5 for a maximum score of 60 if all 12 marbles were fully buried.

#### Social memory

A three-chamber paradigm was used to examine social memory and novelty, measured as preference to spend time with a novel versus familiar mouse ^55^. The test apparatus had 3 equal size chambers (each chamber: length: 36 cm; width: 36 cm; height: 46 cm) with the right and left ones containing an inverted wire mesh cup to hold the stimulus mice and an overhead camera (ThorLabs). On day 1 of the protocol, stimulus mice (same background strain, sex, and age of the experimental mice) were habituated to the testing chamber and to being in the wire cup for 15 min. Experimental mice were habituated to the testing chamber with empty wire cups for 15 min (stimulus mice were not present). On day 2, one stimulus mouse is placed in a cup (the side of the chamber will vary in case there is a side preference); the other cup is empty. The experimental mouse is placed in the center chamber and allowed to explore all chambers freely for 15 min. The amount of time it spends in the chamber with the cup containing the mouse versus the empty cup is recorded as a measure of sociability. This phase of the test also serves to familiarize the experimental mouse to this particular stimulus mouse which will be the “familiar” mouse on test day 3. On day 3 (∼24 hr later), the protocol of day 2 is repeated using the same stimulus mouse so that the experimental mouse can “re-familiarize” itself with this conspecific. Then, 30 min later, the same familiar mouse is placed in the opposite cup and a novel stimulus mouse is placed in the other. During 15 min of exploration, the amount of time that the experimental mouse spends with the novel mouse versus the familiar mouse is recorded as a measure of social memory.

### *In vivo* bioluminescent imaging

Bioluminescent imaging (BLI) was conducted using a Lago X instrument and Aura imaging software v.4.0.7 both from Spectral Instruments Imaging. For *in vivo* imaging, mice were anesthetized with isoflurane (3-4% for induction) then maintained at 1-2% in 100% oxygen (0.8 L/min), injected intraperitoneally with 25 mg/kg Akalumine-HCl (Sigma), and then imaged 5, 10, and 15 min later. Imaging parameters were 30 sec exposure time and 25 × 20 × 15 field of view. The maximum total emission signal was assessed with Aura software.

### Safety switch activation *in vivo*

A separate cohort of mice underwent the immunosuppression and hSTRpc-BDNF transplantation surgery protocols as described above (*n*= 10 mice). Ten days post-transplant, the mice received rapamycin (*n*= 7 mice) or vehicle (*n*= 3 mice) via intraperitoneal (i.p.) injection once daily for 3 days. Rapamycin (LC Laboratories) at 20 mg/kg was dissolved in 4% ethanol, 5% polyethylene glycol 300 (PEG300), 5% Tween-80 in Dulbecco’s PBS, vortexed, and sonicated for 5 min in a water bath. Mice were imaged 2 days post-injection and one of the mice that received vehicle died after imaging. Mice were euthanized 1 or 2 weeks after the last rapamycin injection; brains were collected for histochemical processing.

### Tissue preparation and immunostaining

Five to six weeks after transplantation surgery, mice were deeply anesthetized with sodium pentobarbital (FatalPlus; Med-Vet) and transcardially perfused with saline solution. Brains were removed and, for a subset of them, the striatum was dissected from one hemisphere and was flash frozen and stored at -80°C until use for ELISA. The other hemisphere was immersion-fixed overnight in 4% paraformaldehyde in 0.1M phosphate buffered saline (PBS; pH 7.4), cryoprotected in 30% sucrose/PBS, and sectioned (30 µm, coronal) using a freezing microtome. Free-floating sections were processed for fluorescent immunostaining to visualize the location of the transplanted hPSCs using the human nuclear marker, Ku80 (made in mouse: MAB1281, Sigma; or made in rabbit MA5-14953; ThermoFisher). To investigate the differentiation profile of the transplanted hPSCs, every 8^th^ section of the brain was processed for double-fluorescent immunostaining for Ku80 and one of the following: Microtubule-associated protein 2 (MAP2; 8707, Cell Signaling), doublecortin (ab18723, AbCam), NeuN (ABN78, Sigma), GFAP (Z0334, Dako), or the oligodendroglial marker, O4 (07139, Millpore). To assess the effects of transplanting hPSCs on Htt aggregates or the effects of engrafted cell elimination on inflammation, sections were double-immunolabeled for human nuclei and either Htt (1:200; clone EM48, Millipore) or IBA-1 (1:1,000; WAKO).

### BDNF ELISA with acid-extraction of brain lysates

The striatum from one brain hemisphere was dissected from a subset of the 10-11 week-old male and female R6/2 mice that were transplanted with hSTRpc or hSTRpc-BDNF as well as from R6/2 and WT mice given intra-striatal vehicle injections (n=4-6 mice / group). Tissue was processed for use with the Mature BDNF Rapid^TM^ ELISA Kit: Human, Mouse, Rat (Biosensis, cat#BEK-2211). Acid-treated tissue samples were used since BDNF is typically bound to its receptors and chaperones in many tissues which hinders its detection by ELISA; acid-extraction releases bound BDNF. Therefore, tissue was suspended in acid-extraction buffer [50 mmol/L sodium acetate, 1 mol/L NaCl, 0.1% Triton X100 with glacial acetic acid added until pH 4.0 was achieved] containing a protease inhibitor cocktail (cOmplete mini tablets, Roche cat. no. 11836153001). Homogenates were prepared via probe sonication for 5-7 sec, followed by a 30 min incubation on ice and then another bout of sonication. Next, they were centrifuged (14, 000g) at 4°C for 30 min. Supernatants were collected and neutralized in phosphate buffer pH 7.6. The neutralized sample was used in the ELISA, according to the manufacturer’s instructions. Samples were run in duplicate, the resulting optical densities were averaged and used to interpolate concentrations (ng/ml) using a standard concentration curve.

### Fluoro-JadeB staining

Fluoro-Jade B staining was used to assess if the safety switch was activated with rapamycin resulting in cell death. It is an anionic fluorescein derivative that stains cells that are in the process of degenerating. Staining was performed according to the manufacturer’s instructions (Biosensis, Australia). Briefly, slide-mounted sections were incubated in a basic ethanol solution, 70% ethanol and then water. Background was blocked with a 0.06% potassium permanganate solution for 10 min and then, after washing with water, stained with 0.0004% Fluoro-JadeB in acetic acid for 10 min. Slides were washed, dried cleared with xylene, and coverslipped with DPX.

### Imaging and quantification of immunostaining

Immunofluorescent staining in the striatum was imaged with a Stellaris 5 confocal microscope platform from Leica Microsystems or a Keyence microscope using a 20X or 63X objective (1024 x1024 pixel resolution). For colocalization analysis of Ku80 with NeuN, O4, MAP2, DCX, or GFAP, confocal Z-stack images (20X objective; step size = 0.44 µm; 25 images/stack) were taken of the entire engraftment area (2-5 sections/mouse; 1-12 images/section; n=3-5 mic /group). Images were analyzed with Imaris image analysis software (v.9.9, Oxford Instruments) using the deconvolution and co-localization features. Intranuclear Htt accumulation was evaluated in two confocal Z-stack images (40X objective; step size = 0.4 µm; 25 images/stack; 290 × 290 µm) that were taken adjacent to the engraftment area or to a comparable area in the vehicle mice (2 sections/mouse; 2 images/section; n=5 mice/group). These images were analyzed with ImageJ (1.54g) software using background subtraction, thresholding, and analyze particles functions. If the immunostaining was performed in multiple sets, quantifications were normalized to the WT-veh group of that staining set and cohort. The researchers performing quantitative analyses were blind to the genotype and treatment of the mice.

## Data Availability

The data that support the findings of this study are available from the corresponding author, upon reasonable request.

## Acknowledgements

This work was supported by funding from Taube Philanthropies and the Jean Perkins Foundation to F.M.L. The work in M.H.P.’s laboratory was supported by the Taube Philanthropies, the Laurie Kraus Lacob Fund for Translational Research, and through the Sutardja-Chuk Chair in Definitive and Curative Medicine. S.S is supported by the Hereditary Disease Foundations’ Postdoctoral Researcher Fellowship.

## Author Contributions

D.A.S., S.S., M.P. and F.M.L. conceived and designed the experiments; D.A.S., S.S., T.C., G.C., T.S.C., S.G., T.L.M.M., R.M., and J.S. performed the experiments; D.A.S., S.S., T.C., G.C., and T.S.C. analyzed the data; D.A.S. and S.S. performed the statistical analyses; N.U. and R.M. contributed to the methodology; D.A.S. and S.S. wrote the manuscript and M.P. and F.M.L. critically revised it.

---

**Supplementary Figure 1.**
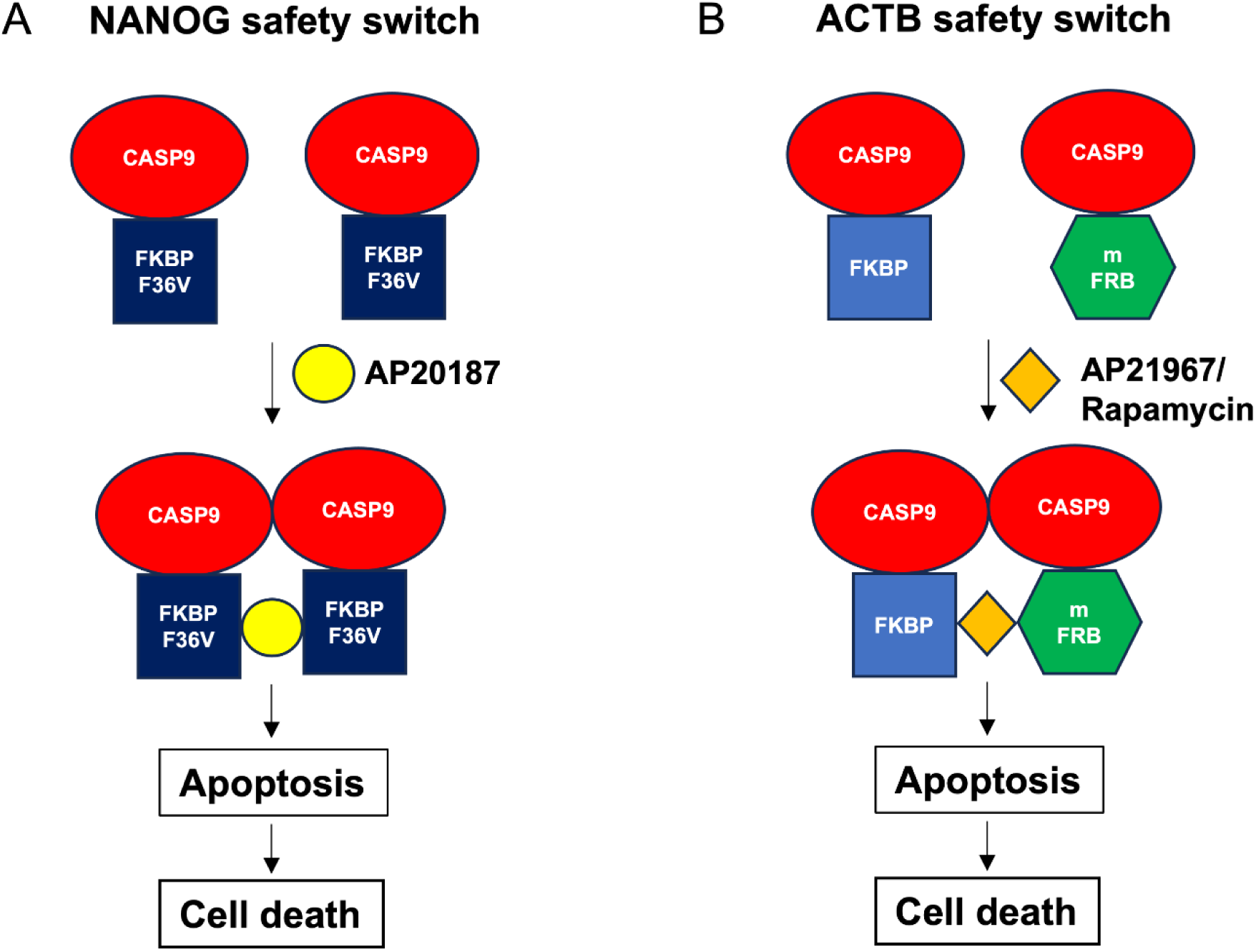
Orthogonal dual safety switches for PSC based therapies. (**A**) Schematic representation of the NANOG safety switch activation with AP20187. The safety switch consists of inducible Caspase 9 (CASP9) knocked into *NANOG* locus and treatment with AP20187 results in formation of CASP9 dimer leading to apoptosis and cell death selectively in pluripotent stem cells. FKBP F36V denotes a mutant FKBP domain. (**B**) Schematic representation of the ACTB safety switch activation with AP21967 or rapamycin. The safety switch consists of orthogonal inducible CASP9 knocked into *ACTB* locus and treatment with AP21967 or rapamycin forms a CASP9 dimer leading to apoptosis and cell death in all cell types. mFRB denotes mutant FRB domain.

**Supplementary Figure 2.**
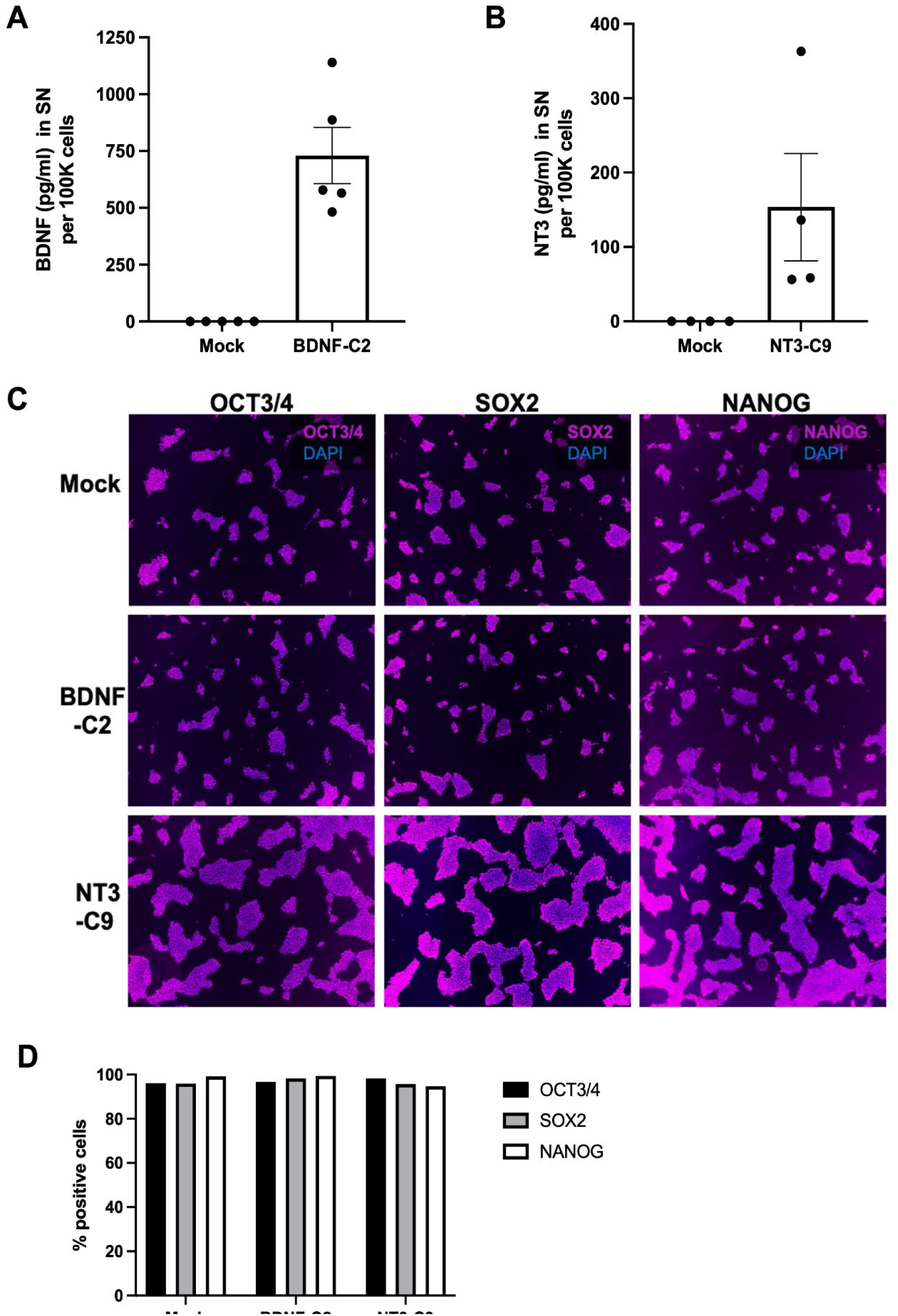
Neurotrophin levels in supernatants and expression of pluripotency markers in BDNF and NT3 overexpressing hPSCs. (**A, B**) Bar graphs showing the quantification of ELISAs for BDNF (**A**) and NT3 (**B**) in the supernatants (SN) of gene targeted single cell ESC clones BDNF-C2 and NT3-C9, respectively. Mock cells were used as a negative control. Data is represented as concentration of BDNF or NT3 (pg/ml) per 100K cells in the SN. (**C**) Representative immunostaining images for pluripotency markers OCT3/4, SOX2 and NANOG in mock, BDNF-C2, and NT3-C9 PSCs. (**D**) Frequency of OCT3/4, SOX2 and NANOG in Mock, BDNF-C2 and NT3-C9 PSCs. Immunostaining images (**C**) were quantified as area of corresponding marker staining relative to area of DAPI staining and represented as percentage of positive cells.

**Supplementary Figure 3.**
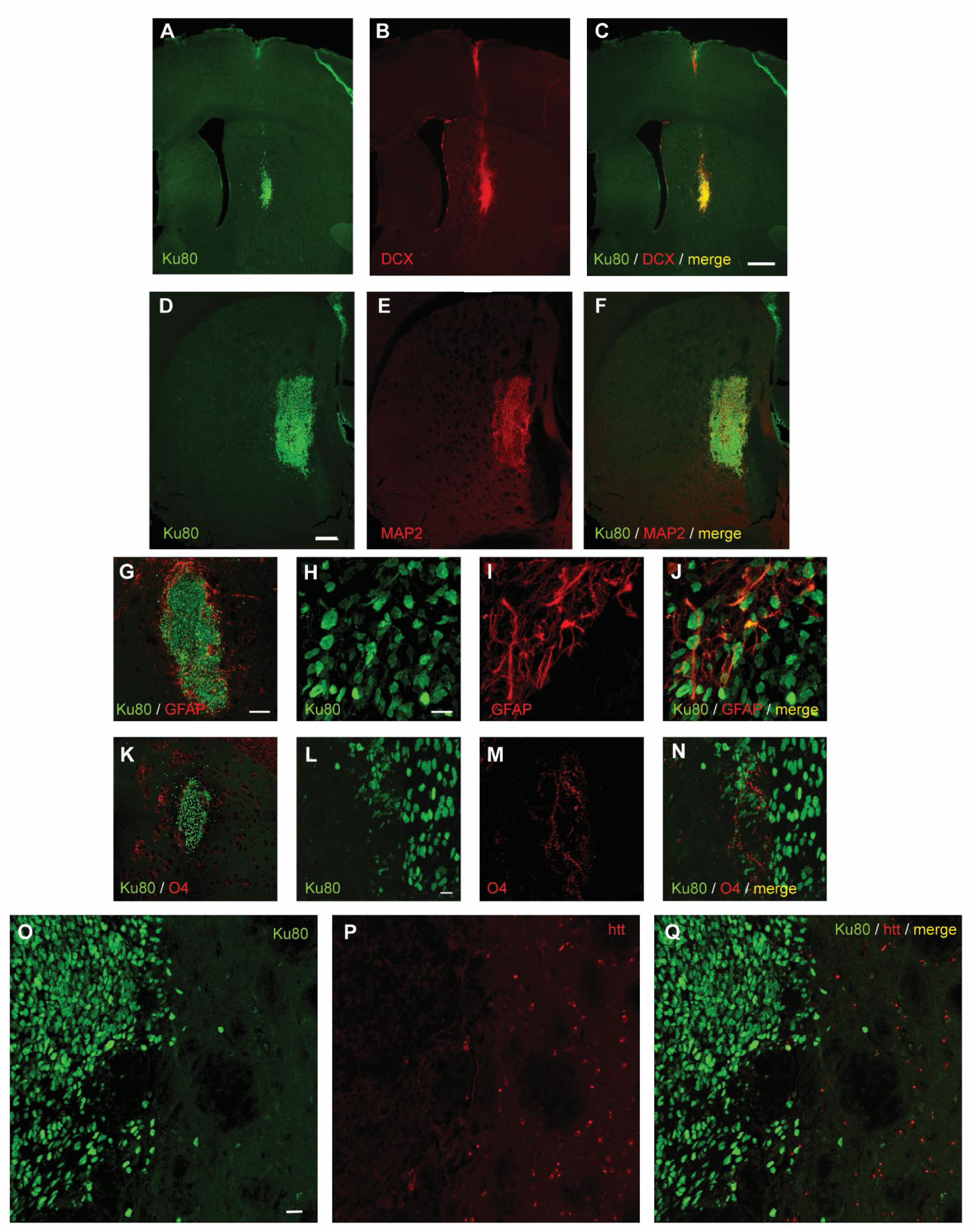
Intrastriatal transplants of hSTRpcs and hSTRpc-BDNF in R6/2 mice survive and colocalize with doublecortin and MAP2 but not huntingtin. (**A-C**) Representative photomicrographs of the STRpc-BDNF engraftment area in the striatum of an R6/2 mouse immunostained for the human nuclei marker, Ku80 (**A**, green), doublecortin (DCX) (**B**, red), and the merged channels (**C**, yellow). Scale bar in C = 300 μm and applies to A-C. (**D-F**) Representative photomicrographs of the hSTRpc engraftment area in the striatum of an R6/2 mouse immunostained for the human nuclei marker, Ku80 (**D**, green), microtubule-associated protein 2 (MAP2) (**E**, red), and the merged channels (**F**, yellow). The engraftment areas were smaller than those seen with hNPCs (see Fig. 4) and the circular cell patterns were not detected in the mice that received hSTRpc or hSTRpc-BDNF transplants. Scale bar in D = 200 μm and applies to D-F. (**G-J**) Immunostaining for human nuclei and GFAP or (**K-N**) oligodendrocytes showing that few of the transplanted cells differentiates into these lineages. Scale bar in G = 100 μm and applies to G and K. Scale bar in H = 10 μm and applies to H -J. Scale bar in L = 10 μm and applies to L-N. (**O-Q**) Representative photomicrographs of the hSTRpc-BDNF engraftment area in the striatum of an R6/2 mouse immunostained for the human nuclei marker, Ku80 (**O**, green), huntingtin (htt, EM48)(**P**, red), and the merged channels (**Q**, yellow) showing a lack of co-localization. Scale bar in O = 10 μm and applies to O-Q.

**Supplementary Figure 4.**
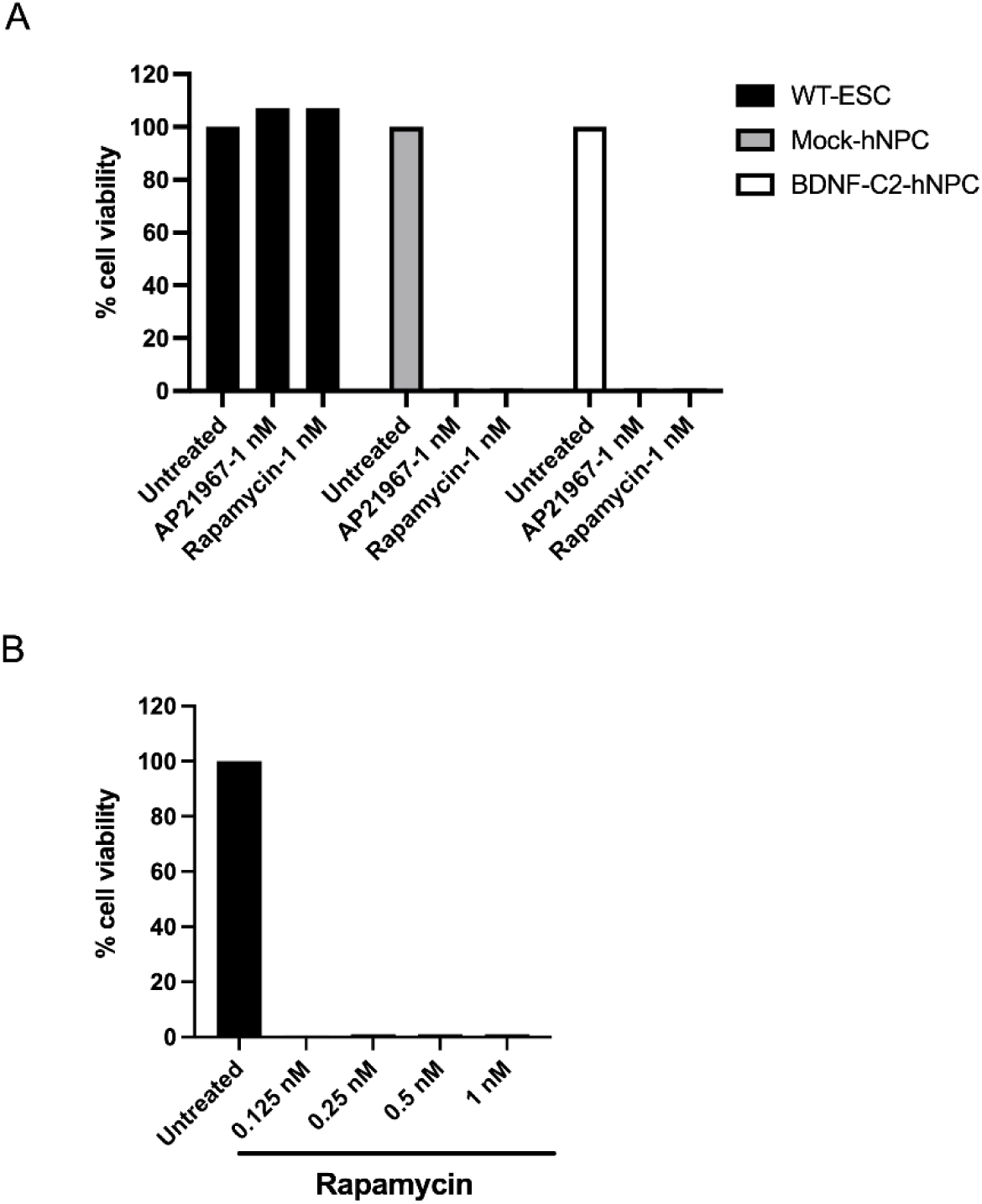
*In vitro* validation of safety switch activation with AP21967 and rapamycin in NPCs and STRpcs. **A.** Cell viability of hNPCs derived from Mock and BDNF-C2 hPSCs post AP21967 or rapamycin treatment (1 nM) for 2 days. WT ESC was used as control. Cell viability was determined through an MTT assay, and the viability is represented as percentage relative to untreated control. **B.** Cell viability of hSTRpcs derived from BDNF-C2 PSC post rapamycin treatment. hSTRpcs were treated with different concentrations of rapamycin, as indicated, for 2 days. Cell viability was determined through an MTT assay, and the viability is represented as percentage relative to untreated control.

